# Liquid crystalline mesophase spacing as a quantitative predictor of release kinetics for co-loaded hydrophilic and hydrophobic payloads

**DOI:** 10.64898/2026.06.26.734853

**Authors:** Sophia R. Dasaro, Shruti S. Sawant, Alexa Stern, Lucas D. Johnson, Caleb F. Fretz, Malinda Salim, Nigel Kirby, Ben J. Boyd, Brian K. Wilson, Gregg Duncan, Qi T. Zhou, Kurt Ristroph

## Abstract

Liquid crystalline mesophases exhibit structurally programmable internal architectures that enable co-loading of chemically orthogonal molecules within a single composite material. Realizing the potential of these materials for drug delivery requires a quantitative understanding of how tuning the composition affects internal mesophase architecture and consequently performance metrics such as payload release. Here, Flash NanoPrecipitation with hydrophobic ion pairing is used to prepare nanocarriers containing liquid crystalline mesophases co-encapsulating two compounds from widely different chemical classes: hydrophilic polymyxin B (logP –6) with one of four hydrophobic co-core materials (logP 7-11), achieving >75% encapsulation efficiency and up to 32% and 50% mass loadings for polymyxin and co-core. Synchrotron SAXS is used to quantify characteristic mesophase repeat spacing, which is found to be tunable as a function of composition. A strong correlation between d-spacing and polymyxin release rate is presented. Co-core chemistry and weight fraction jointly govern mesophase architecture, and repeat distance emerges as a structural metric linking these to the hydrophilic payload release kinetics. Mucus diffusivity and antibacterial efficacy are assessed as independent performance metrics, and results corroborate the release behavior. These findings establish a quantitative framework connecting material composition, mesophase architecture, and functional performance that can be applied toward rational co-formulation design.

ToC Graphic Text
Flash NanoPrecipitation yields liquid crystalline nanocarriers co-encapsulating with high efficiency payloads with widely distinct physicochemical properties. Synchrotron SAXS establishes characteristic repeat spacing as a quantitative structural metric directly governing hydrophilic release kinetics, providing a rational design framework linking mesophase architecture to functional performance across a range of payload structures.

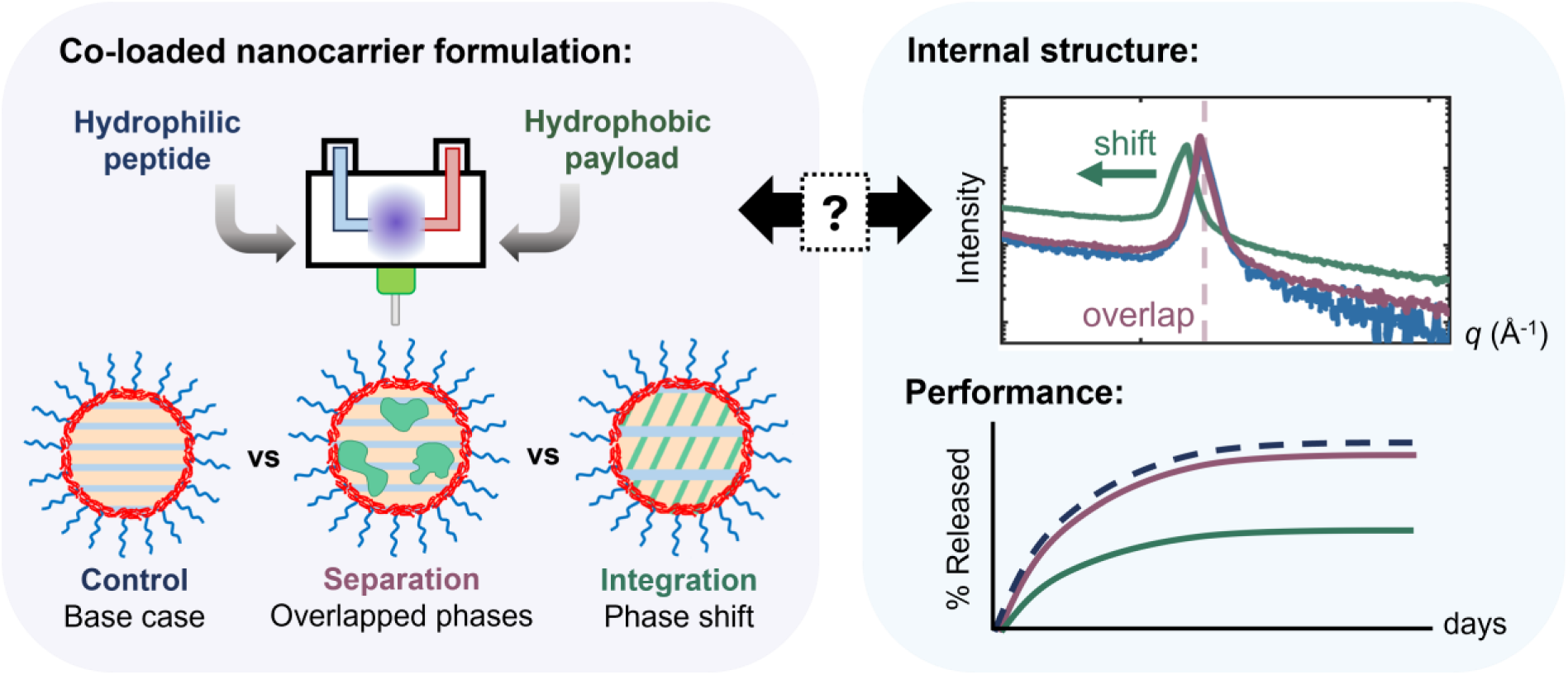

## Introduction

Precision delivery of multiple therapeutics using a single delivery vehicle requires platforms that enable the co-encapsulation of chemically diverse payloads while maintaining independent control over the release rates of each. Combination therapies that rely on drug synergy for therapeutic efficacy represent a compelling clinical need for co-formulation platforms, yet such strategies remain technically challenging as the active pharmaceutical ingredients (APIs) often exhibit conflicting physicochemical characteristics and delivery requirements[1–5]. Consequently, conventional administration of combination therapies relies on coordination across multiple dosage forms, complicating temporal alignment of therapeutic windows and limiting synergistic pharmacological effects[3, 6–8]. A single, tunable delivery platform capable of accommodating multiple payloads with largely different physicochemistries into a single functional composite material would address these limitations and expand the available repertoire of delivery strategies for modern combination therapies.

Liquid crystalline (LC) mesophases are well-suited to serve as such a platform. LC phases exhibit complex internal architectures such as bicontinuous or lamellar networks of hydrophilic aqueous channels and hydrophobic lipid domains that provide spatially distinct environments, enabling co-encapsulation of both hydrophilic and hydrophobic APIs[9–11]. LC phase geometry can be modulated by environmental factors (such as pH) and formulation parameters (such as lipid composition). A goal of this work was to determine design rules for how these parameters control payload release rates, and how tuning composition affects material performance. We focus specifically on API-loaded LC phases nanoencapsulated using a polymer or surfactant stabilizer, an attractive approach for protecting sensitive cargo from harsh physiological environments prior to release [12]. Realizing the full potential of LC nanocarriers (LCNCs) as a co-formulation platform requires a quantitative understanding of how formulation parameters govern mesophase architecture and, in turn, functional performance. This relationship that has not been systematically established for LCNCs co-loading hydrophobic and hydrophilic payloads.

There are two approaches to manufacturing colloidal LC phases: top-down dispersion of a bulk mesophase by applying external energy (e.g., high pressure homogenization or sonication) and bottom-up antisolvent precipitation techniques in which the lipid mixture, API, and stabilizer are co-dissolved in a non-polar solvent then mixed against a polar anti-solvent in a controlled fashion to induce self-assembly[12, 13]. Bottom-up approaches offer distinct advantages in scalability and reproducibility, with turbulent-flow micromixers representing the most robust technology for large-scale manufacture of colloidal LC phases[14–19]. This area remains underexplored outside the context of lipid nanoparticles designed for genetic medicine.

A recent report by Almunif et al. demonstrates the successful production of size-controlled bicontinuous nanospheres (BCNs) formed by amphiphilic block co-polymers via Flash NanoPrecipitation (FNP), a scalable antisolvent precipitation technique that relies on turbulent flow micromixers to evenly distribute formulation components prior to nanocarrier self-assembly[18, 20]. We previously combined FNP with *in situ* hydrophobic ion pairing (HIP) to produce oleic acid-based LCNCs encapsulated within an amphiphilic diblock copolymer; those formulations achieved efficient (>90% for both payloads) co-encapsulation of a hydrophilic antibiotic peptide, polymyxin B (PMB), and model hydrophobic payloads[21, 22]. Those studies established the kinetics of LC phase formation in co-loaded LCNCs and provided manufacturing guidelines for scale-up[21, 23]; however, a detailed evaluation of co-loaded LCNC performance and efficacy – especially controlled release – as a function of formulation parameters, remained unknown.

We here address this gap. LCNC formulations of polymyxin B (PMB) and one of four hydrophobic model APIs, referred to herein as ‘co-cores’, (vitamin E acetate (VEAc), cholesterol, polycaprolactone, or methyl oleate) were prepared and systematically evaluated across a formulation design space spanning co-core chemistry, co-core weight fraction, and HIP charge ratio[24]. Synchrotron SAXS was employed to resolve mesophase ordering and characteristic repeat spacing across three pH conditions, and *in vitro* release kinetics were measured under matched conditions (deionized water, phosphate buffered saline (PBS) at pH 7.4, and PBS adjusted to pH 4.5) to establish direct correlations between mesophase architecture and hydrophilic payload release. The release of co-formulation PMB and VEAc were tracked simultaneously in the presence and absence of a hydrophobic sink for up to 5 days.

Co-core identity, weight fraction, and HIP charge ratio collectively modulate mesophase architecture, with d-spacing emerging as a previously-unknown key structural metric linking formulation parameters to PMB release kinetics. Mucus diffusivity and antibacterial efficacy against *Pseudomonas aeruginosa* were assessed as independent functional performance metrics; trends across formulations corroborate release behavior, providing further validation of the mesophase structural framework. Together, these findings establish a quantitative, mechanistically informed structure-function relationship for co-loaded LCNCs that enables rational formulation design for co-encapsulating hydrophilic and hydrophilic actives together into a single delivery vehicle.

## Main text

### Aqueous environment & co-encapsulation prompt rearrangement of liquid crystalline mesophases

It is well established that pH and ionic strength play important roles in governing liquid crystalline mesophase assembly by influencing the charge state and local area occupied by the headgroup of the amphiphilic molecule(s) utilized to produce LC phases[25–27]. When the headgroups of an amphiphile are charged, the effective headgroup area is comparably larger due to electrostatic repulsions[26]. For amphiphiles with acidic headgroups, at a pH below the pK_a_, the headgroups are neutral and an oil-like behavior is observed. In classical LC systems, this can be exploited to drive phase transitions for controlled release applications. In the LCNC system presented here, there is an added layer of complexity: the state of the hydrophobic ion paired complex. Both ionic strength and pH can compromise the stability of ion paired complexes and disrupt LC phase formation[22, 24]. Therefore, it is important to establish how the LCNCs presented here evolve upon exposure to different aqueous environments (**Figure 1**).

**Figure 1:**
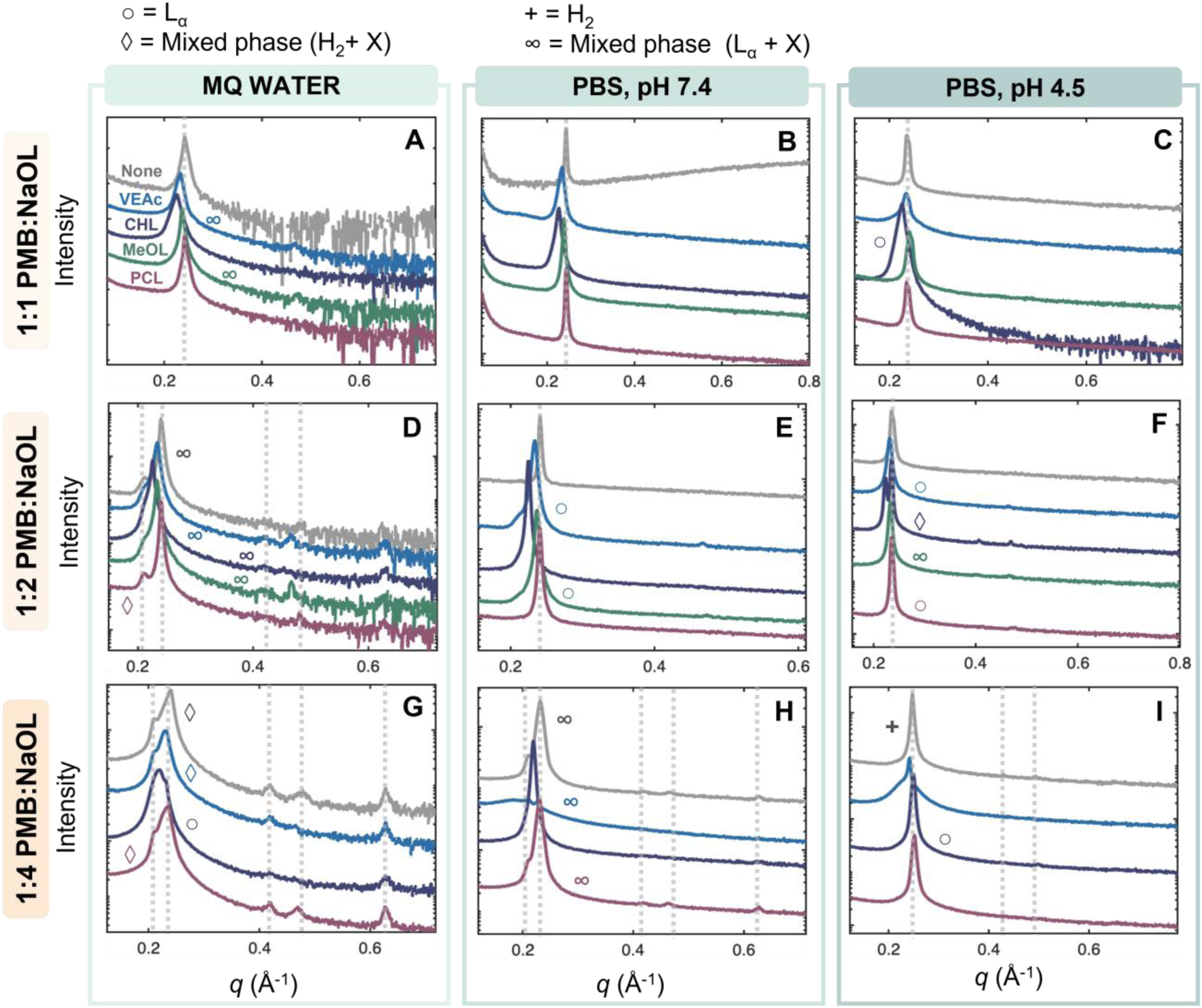
SAXS profiles of liquid crystalline nanocarriers containing 0% or 10% of each hydrophobic co-core at a 1:1 (A-C), 1:2 (D-F), or 1:4 (G-I) charge ratio at three environmental conditions. Co-core type is color coded following the nomenclature in panel A. Where identifiable, LC phases are denoted by symbols: ◊ = Mixed phase where one identified phase is inverse hexagonal (H_2_), + = Inverse hexagonal, ○ = Lamellar (L_α_), and ∞ = Mixed phase where one identified phase is lamellar. Data for 10% MeOL LCNCs at a 1:4 charge ratio was collected for samples in MQ water only as discussed and provided in **Figure S.20**.

Across all formulations, there are two dominant LC phases that comprise the core of the LCNCs: inverse hexagonal (H_2_) and lamellar (L_α_). In some cases, both mesophases may co-exist. However, due to overlapping primary and higher order peaks, deconvolution of the specific phases comprising the LCNC core in this system is challenging to execute definitively, though it has been done before for other surfactant-containing systems[28]. The co-existence of LC phases has been previously reported and, in this work, is rationalized through a discussion of charge ratio[21, 22, 28, 29]. At higher charge ratios (1:2 and 1:4), there is excess charged oleic acid present in the LCNC core, facilitating the assembly of complex internal architectures (H_2_). The co-existence of at least two unique LC phases can subsequently be attributed to a population of LC phases primarily composed of PMB:OL complexes and a subpopulation of LC phases formed primarily by the excess oleate.

Across charge ratios, as ionic strength is increased (transition from MQ to PBS), the intensity of the scattering profiles decreases, complicating the resolution of higher ordered peaks. This effect is likely driven by increased salt presence screening the electrostatic effects that drive the formation of the PMB:OL complexes counterion[25]. In addition to a decrease in overall order, at a pH of 7.4 no pure or mixed H_2_ phases are observed. However, at pH 4.5, oleic acid is primarily neutral, and recovery of more complex internal architectures is observed. The packing parameter (P) is a useful tool in explaining the observed effects[30]. The P relates the relative areas and volumes occupied by the amphiphile head and tail groups to predict LC mesophase formation[30, 31]. At low pH, the P increases, facilitating the formation of structures with negative membrane curvature. In the data, this is reflected by the isolated H_2_ phase present only at a 1:4 charge ratio at low pH.

Comparing the SAXS profiles and primary peak locations for samples with a co-core vs without, clear trends emerge. In all cases, the grey dotted lines (placed to align with the 0% co-core formulations) intersect formulations where PCL_2k_ is the co-encapsulated payload. For all other co-cores, there is a leftward shift of the primary peak toward lower *q*, indicating an increase in lattice spacing. This observation is corroborated by kinetics studies investigating these formulations as well as the tabulated peak indices provided in the SI (**Table S.3, S.4, & S.5**)[21]. Increasing the weight fraction of the hydrophobic co-core beyond 10% did not drive significant changes in the overall LC phase but did induce shifts in the primary peak location further toward the left (**Figure S.1-4**), supporting previous investigations regarding LCNC assembly kinetics and dynamics[21]. To illustrate the modulation of LC phase characteristics by the presence or absence of co-encapsulated payloads, d-spacing analysis is provided in **Figure 2**.

**Figure 2:**
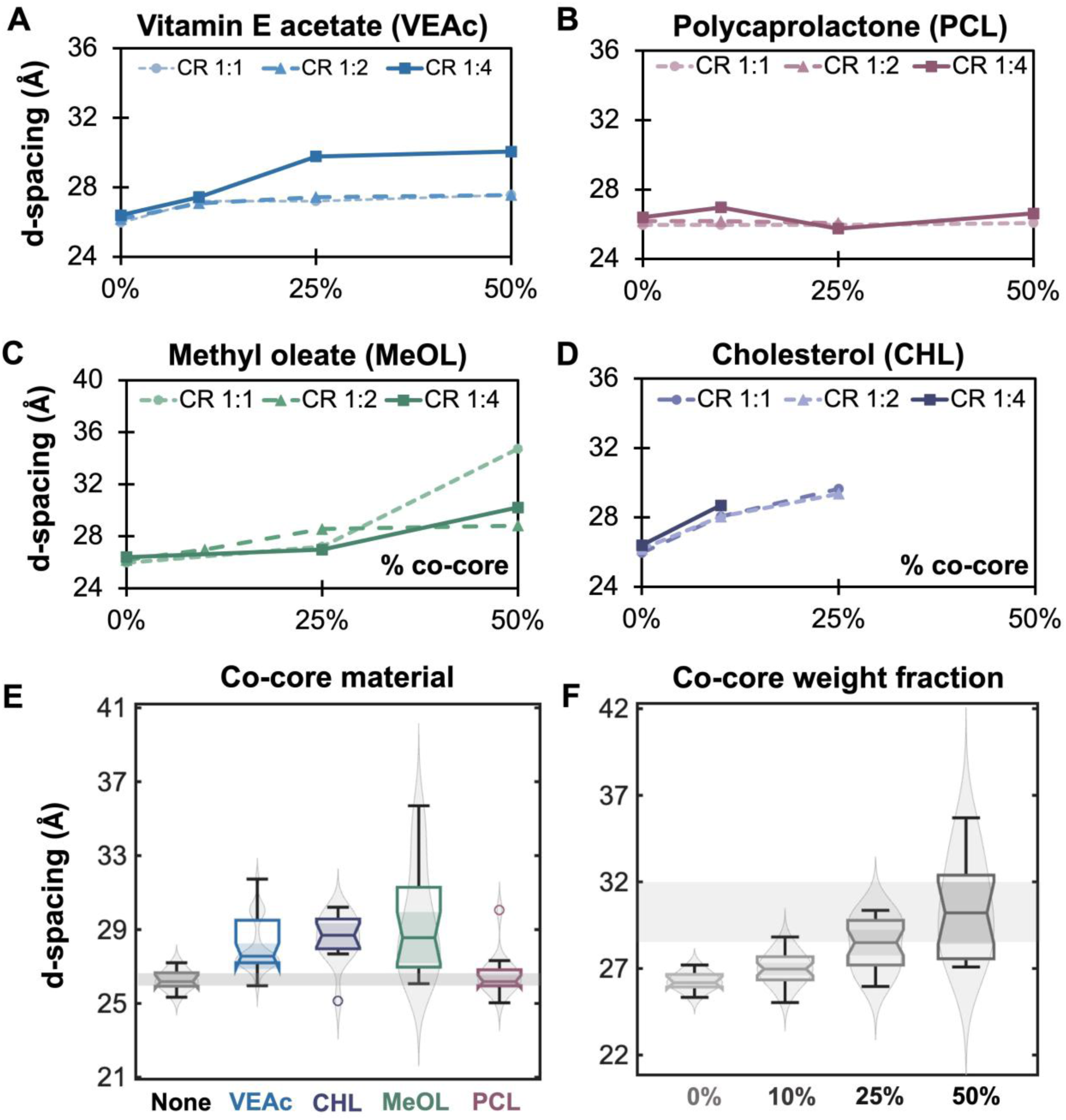
Relationship between d-spacing and co-core weight fraction for all charge ratios and formulations including vitamin E acetate (A), polycaprolactone (B), methyl oleate (C), and cholesterol (D) in MQ water. (E&F) Combined analysis of data in (A-C) to highlight relationships between co-core type and weight fractions where shaded bands are included to visualize important relationships and overlap of shaded regions of the notched bar charts. The d-spacing values were determined using the primary peak location correlated with the most dominant phase (L_α_) determined based on peak intensity.

PCL_2k_ is the only hydrophobic co-core that does not significantly influence the d-spacing spacing of a given LC phase. A proposed hypothesis for this result is that PCL_2k_ phase separates in the nanocarrier core rather than intercalating within the LC phase, as evidenced by the absence of an effect of PCL_2K_ on repeat unit spacing and features that are characteristic of semi-crystalline PCL_2k_ in the LCNC cores (a broad halo or additional feature in the SAXS curves). However, as suggested by **Figure 2 (A, C&D)**, increasing the co-core weight fraction for all other co-cores results in larger d-spacings. This trend was observed across all investigated conditions (**Figure S.5**). The grouped analysis in **Figure 2E** illustrates the significance of these effects where the shaded region of the PCL_2k_ bar overlaps only with the no co-core samples for all charge ratios and weight fractions. While analysis of **Figure 1** shows that charge ratio and environment drive changes in overall LC phase, repeating the grouped analysis presented in **Figure 2** for these parameters reveals that the d-spacing is largely conserved across charge ratios and environment (**Figure S.6**). Essentially, for a fixed LC phase, co-core type and weight fraction will more significantly impact d-spacing compared to adjusting the charge ratio or environment. A decrease in pH in the external environment will disrupt the PMB:OL ion pairs and drive a decrease in overall ordering that does not appear to influence the specific d-spacing of a given LC phase. Together, the analyses performed in **Figure 1&2** identify both environment-driven and formulation-driven effects on LC structure.

### Variations in release media composition inform drug release behavior

A series of initial release kinetics experiments were performed to compliment the SAXS analysis and inform the mechanisms that govern API release kinetics. As discussed, charge ratio, hydrophobic co-core type, and co-core weight fraction all play critical roles in dictating LCNC internal structure and spacing. Therefore, a subset of LCNCs capturing these key variables (6 formulations – 5 from the screen + 1 additional) was selected and analyzed under each of three conditions: PBS pH 4.5 vs. 7.4, and PBS pH 7.4 with vs. without CHAPS. The selected formulations (**Table S.7, Figure S.12 A-D**) compare LCNCs at a 1:1 and 1:4 charge ratio with and without a 25% wt. fraction VEAc or PCL_2k_ co-core. VEAc was selected as a model co-core that induces a shift in the primary peak while PCL_2k_ was selected as it is the only co-core that does not significantly influence d-spacing. The release conditions investigated here isolate effects that are traditionally coupled: ionic strength and hydrophobic sink.

The characterization of the model formulations is presented in **Figure S.12 A-D.** Characterization reveals that an increase in charge ratio correlates with an increase in encapsulation efficiency and decrease in nanocarrier size[24, 32]. At the stoichiometric ratio equivalence point across all co-cores, the nanocarrier surface charge is predominantly neutral (9.10 ± 2.19 mV) while at a 1:4 charge ratio the surface charge is slightly negative (–12.9 ± 1.84 mV) excess counterion partitioning to the particle surface (**SI Section 6.2**). Although the ion-paired complex is predicted to have similar hydrophobicity at 1:1 and 1:4 charge ratios, excess charged oleate at higher ratios likely increases electrostatic repulsion between complexes. This inhibits nuclei coalescence, stabilizing smaller nuclei for longer timescales and enabling polymer adsorption prior to further growth, ultimately producing smaller nanocarriers.[33].

Figure 3 shows the release kinetics data for the model formulations in each of three different media. Pair-wise comparisons of the same formulation in each media type, all else constant, inform the mechanisms that dictate release of the HIP complex from the LCNC. At a fixed pH of 7.4 in presence of CHAPS, the release of PMB from the LCNC is complete in ∼24 hrs, whereas without CHAPS, release is <10% complete across all formulations in that same time frame (Figure 3 A-C**, SI S.14 A-C**). Therefore, CHAPS acts as an effective hydrophobic sink, aiding in the partitioning of the HIP complexes out of the LCNC core, through the PCL_5k_ portion of the nanocarrier shell, and into the bulk media[34]. For each set of curves, the similarity factor (***f*_2_**) was calculated to numerically determine whether the observed difference between the curves was significant[35, 36]. Comparing release with vs. without CHAPS, all computed similarity factor values were less than or equal to 50 (Figure 3 A-C), indicating release profiles were different and that CHAPS significantly influenced the release kinetics of PMB[35, 36]. A similar effect is observed for the release kinetics in the absence of CHAPS, at pH 7.4 vs pH 4.5 (Figure 3 D-F). At pH 4.5, all formulations exhibit release rates > 50% after 24hr while at pH 7.4, none of the formulations achieved a total drug release > 25%. These findings are consistent with the SAXS data, which show that at low pH the primary peak intensity decreases and higher-order peaks are less resolved, suggesting pH induced minimization of PMB:OL electrostatic interactions facilitating faster drug release. Across all media types, the release rate of PMB from the LCNC is slower at higher charge ratios (Figure 3 G-I). This is consistent with historical release kinetics data for the PMB:OL system and expectations for HIP systems[22, 37, 38]. It is also important to note that the release hierarchy presented in Figure 3 and discussed above is conserved across all other pairwise comparisons, which are presented in the SI (**Figure S.14**).

**Figure 3:**
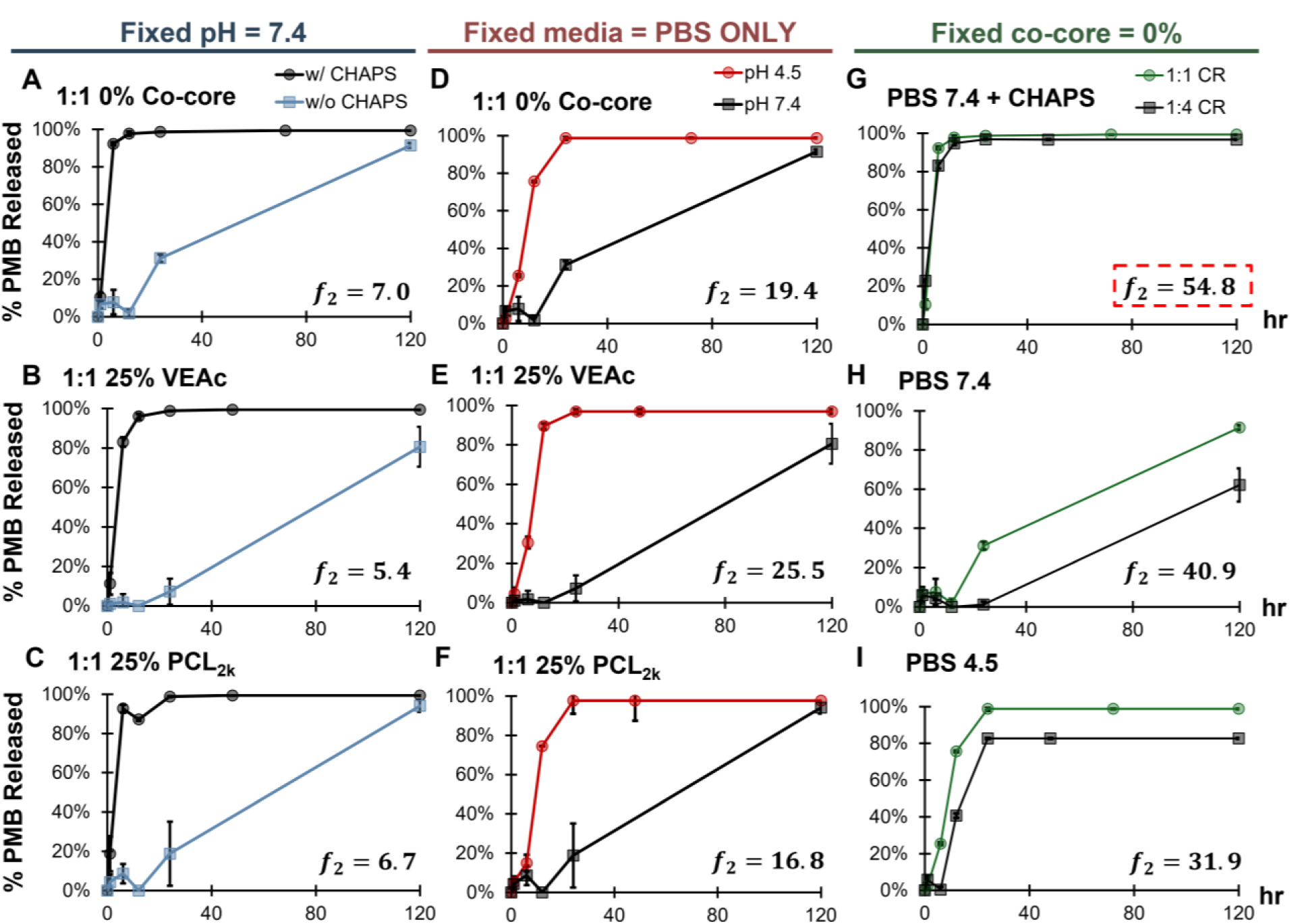
Pairwise comparisons at a fixed pH comparing media containing CHAPS vs without (A-C), in the absence of CHAPS comparing pH (D-E), and at all three media conditions but fixed co-core (CC) (G-I). Error bars represent the standard deviations for *n* = 3 separate tests (some error bars are smaller than the marker size). Panels (A-F) at a 1:4 charge ratio and (G-I) for all other co-cores are presented in the SI and recapitulate the same trends. ***f*_2_** values were determined using the methodology described in the **SI**, **Section 6.3[35, 36].** A dotted box was included in panel G to indicate the only set of curves that are statistically indistinguishable from one another according to the ***f*_2_** analysis.

### Nanocarriers exhibit controlled release compared to free drug

In dialysis-based release experiments, the dialysis membrane itself can act as a barrier to release of the therapeutic necessitating evaluation of the LCNCs with respect to the free drug. For this comparison, the six key formulations and the media in which release rate was the greatest were selected (PBS pH 7.4 with CHAPS). To accurately compare each formulation, a more sensitive timepoint-by-timepoint statistical analysis was performed (**SI, Section 6.4**). The results are presented in Figure 4.

**Figure 4:**
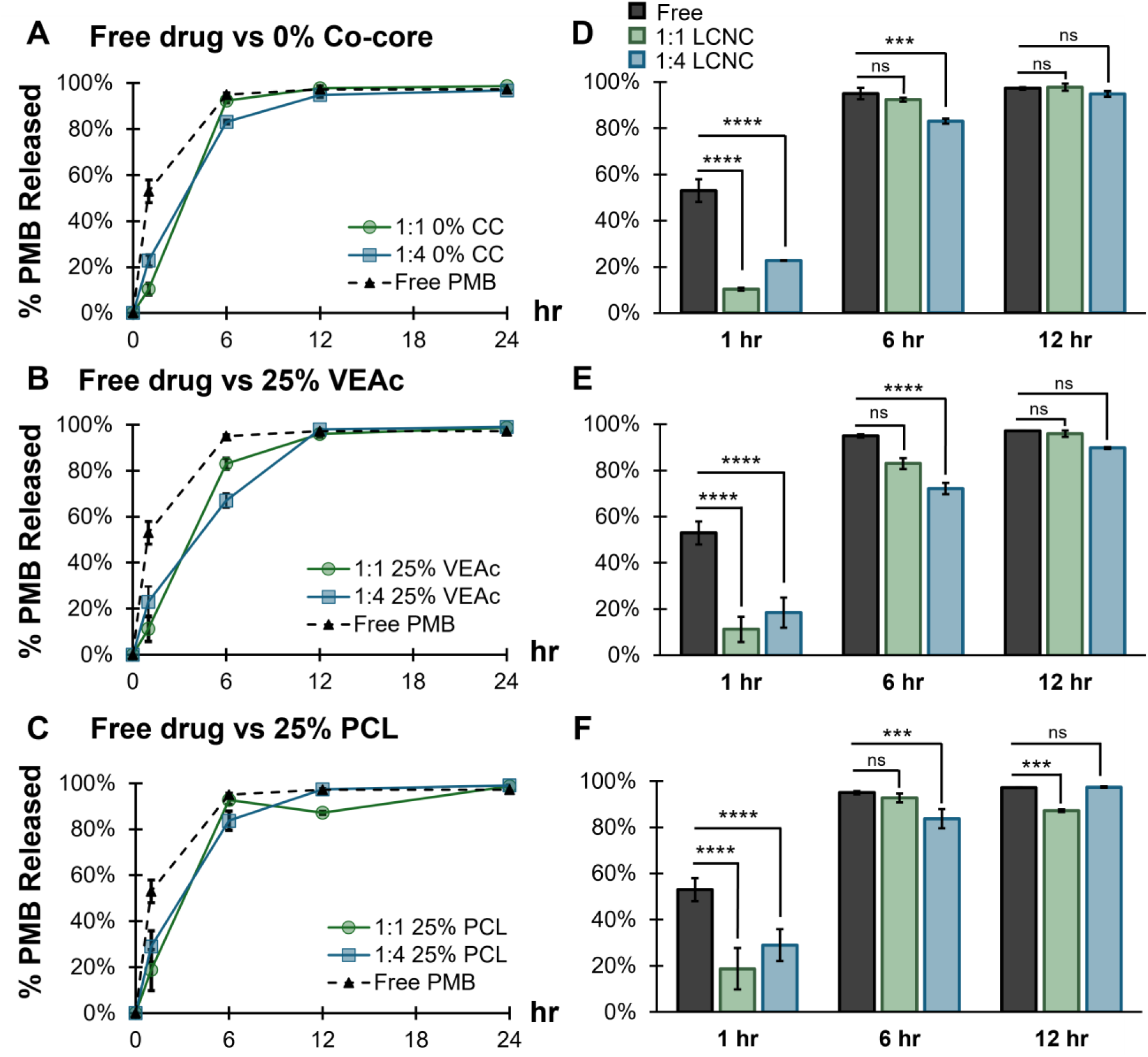
(A-C) Release kinetics profiles of LCNCs without a co-core (CC) (A), with a 25% VEAc co core (B), and a 25% PCL co-core (C) at a 1:1 and 1:4 charge ratio versus the free drug in PBS pH 7.4 with CHAPS. (D-F) Bar charts comparing the average percentage of PMB release at 1 hr, 6 hr, and 12 hr. Statistically significant differences were determined using a two-way ANOVA with Tukey multiple comparisons (**SI Section 6.4**). The significance is denoted by asterisks following: p < 0.0001 = ****, p < 0.001 = ***, p < 0.01 = **, p < 0.05 = *. Error bars plotted as the standard error of all replicates (*n* = 3), and some error bars are smaller than the marker size). The free drug curve in panels A is replicated in panels B&C for ease of comparison.

After one hour, all formulations exhibit a significant difference in the release of PMB compared to the free drug. After 12 hours, only the 1:4 25% VEAc formulation and 1:1 25% PCL_2k_ formulations appear to be significant. Comparing the six-hour time point in Figure 4 D-F reveals that VEAc is a more effective co-core in controlling the release of PMB compared to PCL_2k_. These data suggest that the release kinetics of formulations without a co-core (0% CC) and formulations with a PCL_2k_ co-core (25% PCL_2k_) are the same. Coupled with the SAXS data, this trend provides potential insight regarding the localization of these materials in the LCNC core; namely, that the PCL phase separates form the PMB:OL within the LCNC core in isolated domains, which would explain why the d-spacing was unchanged. Conversely, the observed shift in the primary peak paired with an alteration in release kinetics observed for VEAc suggests that VEAc may be integrating into the PMB:OL LC mesophase, swelling the bilayer, which corresponds with a delay in the release of PMB. Predicting whether a material will phase separate (and conserve the release kinetics of the ion pair) or integrate within an LC phase (and modulate release kinetics) would be advantageous in engineering LCNCs for controlled drug delivery.

### Formulation parameters dictate *in vitro* release kinetics

To explore the parameters that are critical to informing such predictions, a series of formulations co-encapsulating PMB and each of the four hydrophobic co-cores at two weight fractions were subjected to release testing in PBS pH 7.4 with CHAPS. This condition was isolated for physiological relevance as both the blood and the lung contain hydrophobic sinks[39, 40]. These regions are mentioned specifically as a potential application space for polymyxin B-loaded nanocarriers is treatment of severe bacterial lung infections via inhalation or intravenous injections. The overall trends regarding LCNC surface charge and EE were recapitulated by the formulations in this larger screen and are presented in the SI (**Section S6.1**). This observation is expected as the dominant parameter governing these physicochemical properties, charge ratio, is held constant in this screen at 1:4 PMB:OL. The full release kinetics profiles and applied statistical analyses are presented in Figure 5 and discussed in the SI (**Section S6.4 & S6.8**).

**Figure 5:**
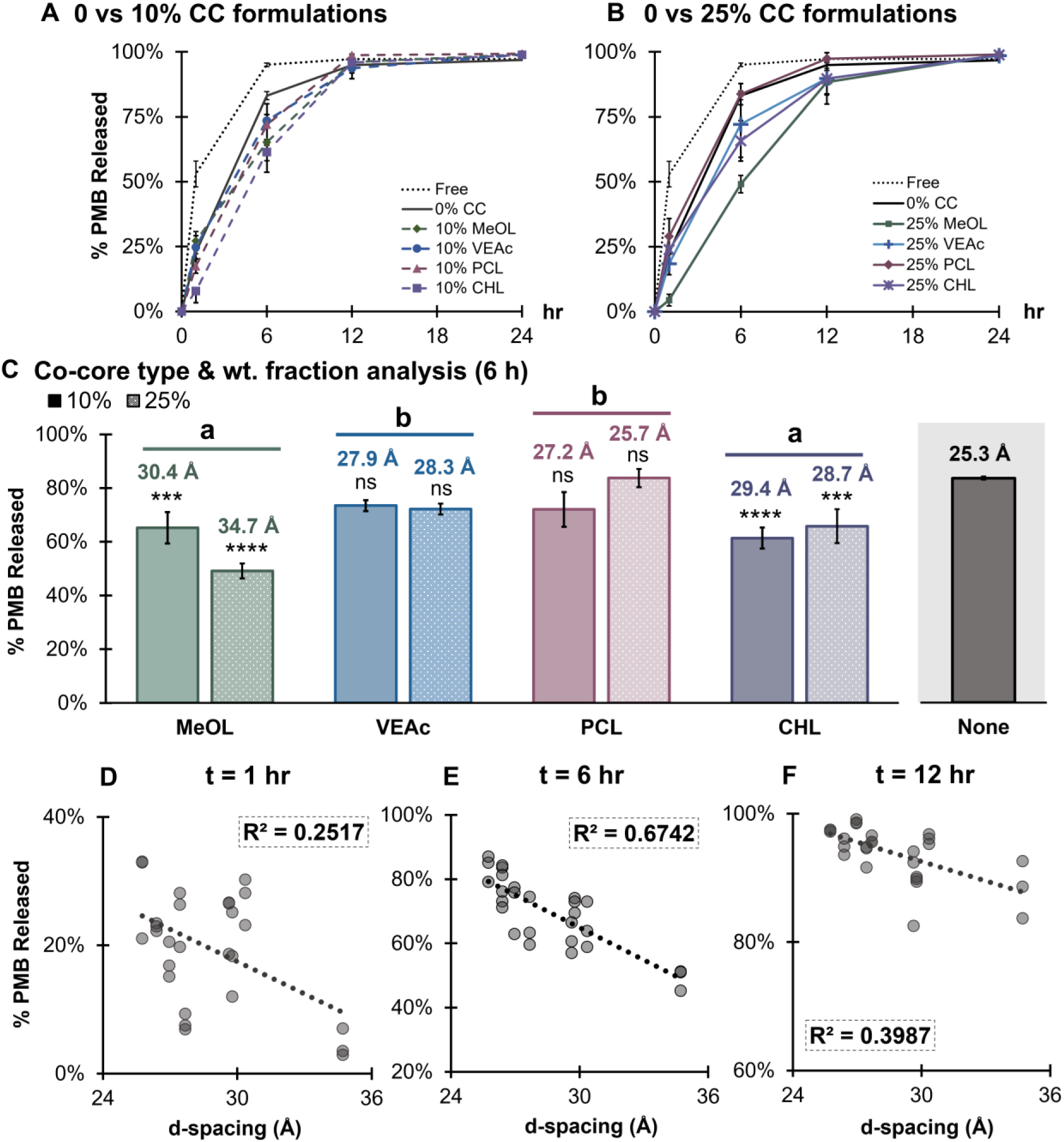
(A & B) Release kinetics profiles for all formulations and free drug controls in the co-core study separated by weight fraction: 10% (A), 25% (B) at a fixed charge ratio of 1:4 in PBS pH 7.4 with CHAPS. (C) Bar charts representing the mean drug release at 6 hr grouped by co-core (CC) type comparing 25% to 10% co-core. The compact letter display represents the combined effect of co-core type and asterisks represent the significance of comparing a fixed formulation to the no co-core sample (isolated grey bar). The statistical analysis in (C) is described in **Section 6.7** of the SI and the d-spacings correlate with values for that sample after exposure to PBS pH 7.4. (D-F) Linear regression line fitting a scatter plot of release percentage vs d-spacing in MQ water (t = 0 hr) across all data points collected. In all plots, error bars represent the standard deviation of *n* = 3 release tests. Asterisks, where present, follow: p < 0.0001 = ****, p < 0.001 = ***, p < 0.01 = **, p < 0.05 = *. The heatmap gradient reflects the magnitude of the p-value where darker shades correlate with smaller p-values. Error bars plotted as the standard error of all replicates (*n* = 3), and some error bars are smaller than the marker size.

All formulations exhibit complete PMB release after 24 hours in PBS at pH 7.4 in the presence of CHAPS. Therefore, significant differences in formulation release behavior were determined by isolating each time point and performing pair-wise analyses. First, all formulations were compared to the free drug to establish a baseline for controlled release (**Table S.11**). At t = 6 hr, formulations exhibit variable release behavior that is (1) co-core dependent and (2) correlated with the determined d-spacing in PBS at pH 7.4. Specifically, samples with d-spacings much larger than the formulations with no co-core (d = 25.3 Å) exhibited release that is significantly slower than the diffusion of the free drug (**SI, Table S.11**). The formulation with no co-core was also compared to the free drug, and the difference in release was found to be insignificant at time points > 1 hr (**SI, Table S.11**). This is also true for the 25% PCL_2k_ samples. Collectively, the 25% PCL_2k_ and no co-core treatments exhibit similar d-spacing, overlapping SAXS patterns, and release that is indistinguishable from the free drug after 1 hr.

While establishing trends in release kinetics relative to the free drug provides a threshold for determining whether PMB transport across the dialysis membrane is sufficiently attenuated by the nanocarrier, pairwise comparisons of each formulation identify which formulation parameters are primarily responsible for driving the observed effects (Figure 5 **C**). Compared to the no co-core control, only methyl oleate and cholesterol result in release kinetics that differ significantly from the other formulations at all time points (**Table S.12**). This is true at both weight fractions; however, the effects are more pronounced for formulations containing a higher co-core weight fraction. These observations correlate with the differences in d-spacing: at a 10% weight fraction, the d-spacing for methyl oleate is 30.4 Å vs 34.7 Å at 25 wt%. Though the absolute difference between d-spacing between each weight fraction is smaller for cholesterol (28.7 Å vs 29.4 Å), the correlated decrease in release kinetics remains the same. We hypothesize that methyl oleate induces the largest increase in d-spacing because of its strong structural compatibility with sodium oleate. As the neutral methyl ester analogue of sodium oleate, methyl oleate can readily intercalate into LC domains through favorable hydrocarbon chain interactions, but lack the charged headgroup that contributes to LC phase organization. Increasing methyl oleate concentration therefore expands the lattice spacing by promoting intercalation and reducing packing efficiency within the ordered structure[41]. VEAc also follows the observed trend where an increase in d-spacing is paired with a decrease in release kinetics, though these effects are not statistically significant compared to the no co-core control and the other co-loaded formulations. These results suggest that altering the co-core material itself has a larger effect on release behavior than varying the weight fraction of a fixed co-core material.

For formulations containing VEAc, the release rate of VEAc from LCNCs was measured and observed to be a function of both charge ratio and weight fraction, in line with observations for PMB. However, statistical analysis reveals these observations are not significant, suggesting that changes in d-spacing play a more significant role in modulating PMB release than the release of the hydrophobic payload (**SI Section 6.5**). The preliminary data also suggest that tuning the release of the hydrophobic co-encapsulated payload by way of mesophase modulation is less straightforward, and that adjusting the payload weight fraction may be a more viable avenue for driving changes in system behavior. To draw such conclusions definitively, further work is necessary applying this model system to a larger suite of hydrophobic APIs with known or predictable release behavior.

### Proposed mechanism for co-encapsulated payload driven differences in release kinetics

In addition to d-spacing, the released fraction of PMB at each time point was compared to the predicted logP values of each material and a rudimentary assessment of thermodynamic compatibility between each co-core and the hydrophobic counterion by applying the Hansen solubility parameters to determine the Hansen distance (**SI, Section 6.6, Table S.8**)[42–44]. The apparent time-dependent nature of the correlation between d-spacing and release behavior may be attributable, in part, to the release media selected for the screen (Figure 5D-F). In the presence of a hydrophobic sink, as discussed above, the release kinetics of all formulations are accelerated, which can make the differences between individual formulations more difficult to resolve. No single analysis nor combination of analyses clearly captured the findings of this study as well as d-spacing alone (Figure 5 D-F). This is in line with work by Martiel *et al.* investigating how different oil modifiers influence liquid crystalline mesophase structure and the release of three hydrophilic actives[45]. That study established that there was no one parameter responsible for clearly summarizing their findings and proposed that their observations were due to a combination of drug hydrophilicity, mesophase complexity, and effect of the oil modifier on membrane fluidity[45]. Considering this, the correlation of d-spacing with LCNC co-core physicochemistry was further evaluated during which it was found that multiple materials with logP values close to that of the counterion exhibited drastically different release kinetics. Similarly, materials predicted to interact poorly with the hydrophobic counterion, determined via the Hansen distance, exhibited different release profiles. For example, cholesterol and PCL_2k_ both had calculated Hansen distances 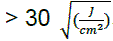, yet only formulations containing PCL_2k_ did not exhibit shifts in the primary peak and modified release fractions. Based on the evidence collected herein, we propose that the observed relationship between d-spacing and therapeutic release may be explained by the mechanism proposed in Figure 6.

**Figure 6:**
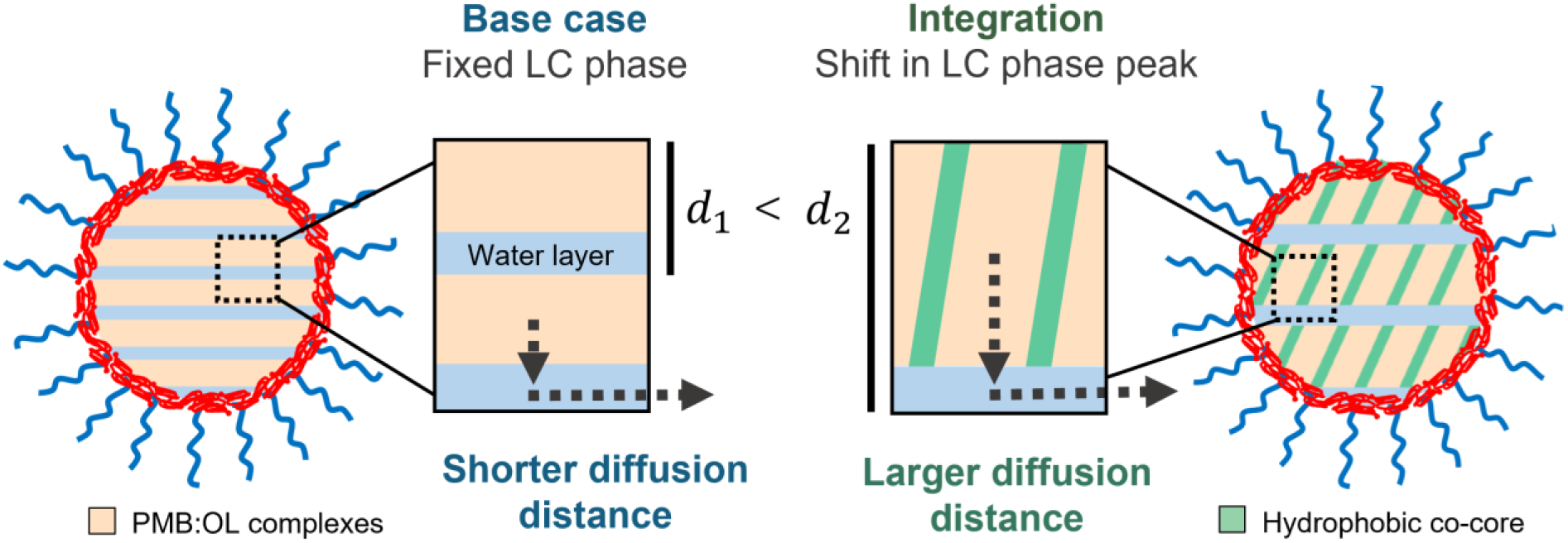
Proposed mechanism for d-spacing driven differences in release kinetics with dashed arrows representing the average distance an HIP complex in the center of the hydrophobic domain of the liquid crystalline mesophase (LC) would have to diffuse to reach a water channel.

While the SAXS data as collected do not provide the required parameters to facilitate an analysis decoupling the water layer from the bilayer, there are still significant differences in d-spacing between formulations. Notably, inclusion of a specific payload did not drive significant changes in overall LC phase architecture, only the spacing of the repeat unit. Thinking of ‘half’ the combined distance of the hydrophobic and hydrophilic domain (d-spacing) as related to the average distance of hydrophobic domain that the hydrophilic payload must diffuse through may explain why samples with higher d-spacing (a longer hydrophobic distance for the payload to diffuse through) release more slowly from the nanocarrier. Since the materials that impact d-spacing are hydrophobic, it can be further rationalized that the hydrophobic co-cores are localized within the hydrophobic domains of the LC phase. This hypothesis agrees with the proposed phase separation of PCL_2k_; formulations containing PCL_2k_ exhibited similar d-spacings and PMB release when compared to the no co-core control.

Ultimately, experimental studies probing these domains and more clearly elucidating this co-localization (or lack thereof) would be necessary to validate these proposed mechanisms. It is also important to note the thermodynamic assessment performed to calculate the Hansen distance is limited by the power of the applied model which is weakened by assumptions made regarding the drug:counterion system. This mechanistic model is based on the more dominant LC phase in this system, L_α,_ and does not explicitly consider how the co-existence of the H_2_ phase impacts the model. Predictive capabilities given the structures of the desired hydrophilic API, hydrophobic API, and hydrophobic counterion could be enabled through a focused effort on advanced modeling techniques and examining how to account for the interaction between the hydrophobic co-core and entire HIP complex.

### Liquid crystalline nanocarrier physicochemical properties impact *in vitro* efficacy against a pathogen

*In vitro* efficacy studies were designed to independently evaluate (i) the influence of charge ratio and (ii) the influence of the hydrophobic co-core at a fixed charge ratio on antimicrobial activity. Based on the release kinetics data, greater differences in performance were expected across formulations with varying charge ratios than among formulations containing different hydrophobic co-cores at a fixed charge ratio. The results of these studies are presented in Figure 7.

**Figure 7:**
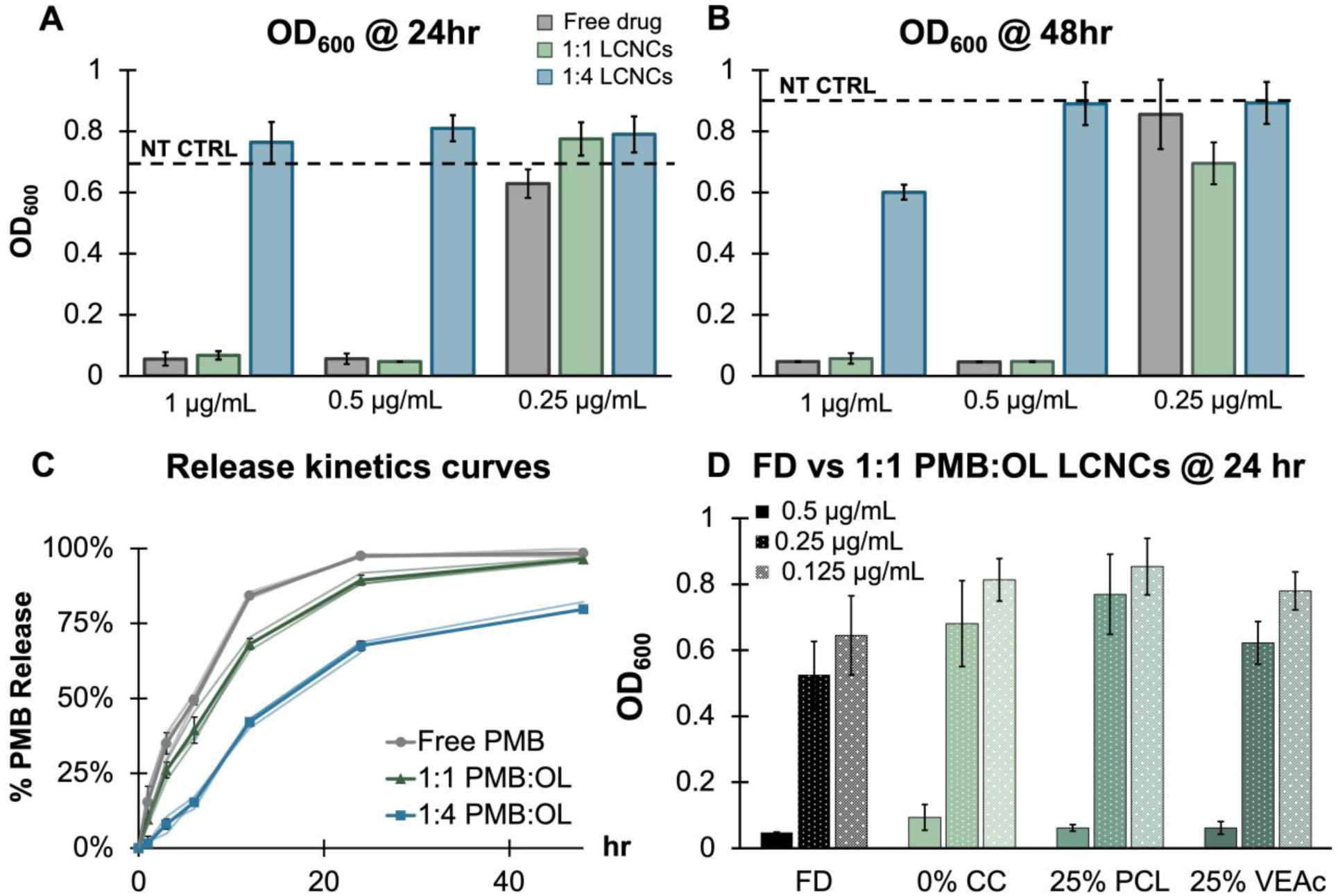
Optical density (OD_600_) for different nanocarrier formulations comparing charge ratio PMB:OL (polymyxin B:sodium oleate) (A&B) and co-core (D). (C) Release kinetics curves for nanocarrier (LCNCs) formulations compared to free drug (FD) in PBS with a nominal pH of 4.5 (160 mM PBS, 0.0045M HCl) and evaluated according to the protocol described in **Section S4.3**. In panel C, the lighter lines are representative of individual replicates. Error bars represent the standard deviation of 12 replicate wells (*n* = 12), and the dashed line represents the OD_600_ for bacteria dosed with an equivalent volume of PBS containing no antibiotic. NT CTRL = no treatment control representative of only bacteria + media. Release data error bars represent the standard deviation of *n* = 3 separate tests. Abbreviations in panel D are CC = co-core, PCL = 2k polycaprolactone, VEAc = vitamin E acetate.

As shown in Figure 7A and B, the 1:1 PMB:OL LCNCs demonstrated greater bactericidal activity over the experimental time course compared to the 1:4 PMB:OL formulations. Optical density at 600 nm was also assessed at 48 h to evaluate bacterial regrowth; however, extending the time course did not reveal additional differences in formulation performance (Figure 7B). This observed difference in efficacy is consistent with the release kinetics data: PMB release proceeds more slowly at a 1:4 charge ratio relative to a 1:1 charge ratio. The alignment between antimicrobial activity and independently measured release profiles further supports the validity of the release testing methodology developed in this work. An additional strength of designing release protocols that mimic relevant biological environments is the ability to evaluate formulation behavior in controlled, less complex media that are compatible with in-line analytical techniques. Specifically, the bacterial growth media utilized herein contain components that can interfere with or compromise analytical instrumentation. By recreating key features of the biological environment in a simplified system, release behavior can be accurately modeled while preserving instrumentation integrity and enabling more robust analytical characterization. In agreement with the proposed hypothesis, Figure 7D indicates that inclusion of a co-encapsulated hydrophobic payload does not measurably alter antimicrobial performance relative to the no co-core control at the same charge ratio for payloads that do not significantly alter release (Figure 7D, **SI Figure S.19B**).

Importantly, the reduced *in vitro* efficacy observed under these conditions does not preclude therapeutic potential *in vivo*. Markwalter et al. previously demonstrated that both 1:1 and 1:4 PMB:OL formulations significantly reduced bacterial colonization in a murine lung infection model [46]. One likely explanation for the discrepancy between the present *in vitro* assay and prior *in vivo* findings relates to differences in available hydrophobic sink conditions. The plate-based growth medium contains limited hydrophobic sink capacity; while proteins are present, they function as relatively weak sinks. Further addition of a stronger hydrophobic sink (e.g., CHAPS) resulted in bactericidal effects and was determined to be non-viable. Therefore, the final aqueous environment was a mixture of the BHI media and the PBS media with a nominal pH of 4.5, ionic strength of 160 mM, and HCl concentration of 0.0045 M. This environment is a contrast to the physiological environments *in vivo* which contain multiple proteinaceous hydrophobic sinks, including serum albumin in the bloodstream and endogenous phospholipids within pulmonary tissue, which may facilitate drug partitioning and release[39, 40]. Furthermore, given the intended application of such therapeutic nanocarriers, the controlled release afforded by the increased charge ratio would allow time for the nanocarriers to diffuse through mucus to the site of the bacterial lung infection without prematurely releasing too much cargo.

### 3.7 Nanocarrier size and surface chemistry modulate mucus diffusivity

The modular approach to nanocarrier formulation afforded by FNP allows for the manufacture of different nanoparticle surface chemistries for a fixed core chemistry. This has been previously explored with respect to mucus diffusivity through selecting different block-copolymer stabilizers to modulate surface charge with neutral particles exhibiting superior mucus-diffusivity compared to those that are strongly negative[47]. Here, all nanocarrier samples have the same block co-polymer stabilizer yet exhibit charge ratio-dependent differences in zeta potential [47, 48]. Therefore, the mucus diffusivity of LCNCs at a 1:1 and 1:4 charge ratio and with no co-core was evaluated to determine if the differences in surface charge quantified here yield new trends. These formulations were isolated because co-encapsulation does not impact the surface charge of the LCNC, and therefore these two formulations captured the suite of surface chemistries present across all the main formulations. The results of this analysis are provided in Figure 8. Following the analysis in work by Lu *et al.,* for particles to be classified as mucus-penetrating, the log_10_MSD_1s_ must be greater than or equal to 0 at a timescale of 1 s[48]. From Figure 8B, LCNCs formulated at a 1:4 charge ratio exhibit an average log_10_MSD_1s_ of ∼ 1 (0.99), indicating strong mucus diffusivity, but LCNCs at a 1:1 charge ratio exhibited non-mucodiffusive behavior. This is significant for two reasons. First, this is the first report of mucodiffusivity for an FNP nanocarrier with a HIP core and shows that the presence of excess counterion does not compromise the efficacy of the dense PEG layer that can be installed on nanocarrier surfaces using FNP. This observation is significant as some of the excess counterion may be expected to arrange near the NC surface and contribute to the observed change in zeta potential away from fully neutral, which could lead to muco-adhesion. Second, it highlights that the difference in nanocarrier size as a function of charge ratio may be a more important factor than zeta potential. LCNCs formulated without a co-core at a 1:1 charge ratio are > 100 nm larger than those formulated at 1:4; despite the zeta potential being closer to zero at 1:1, the smaller NCs at 1:4 diffuse faster.

**Figure 8:**
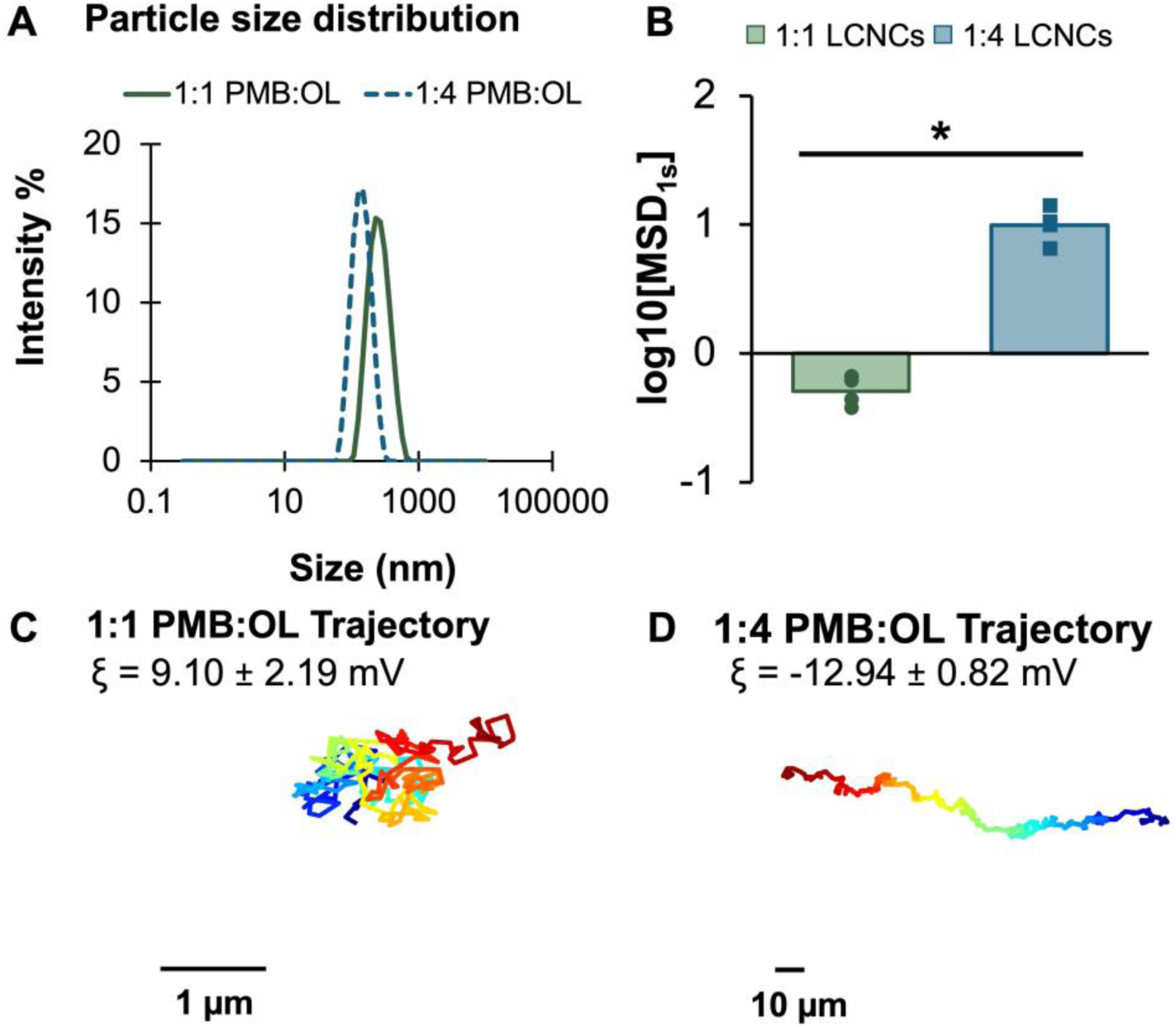
(A) Comparison of size distributions and (B) of log_10_[MSD_1s_] for liquid crystalline nanocarriers (LCNCs) formulated at a 1:1 vs a 1:4 PMB:OL (polymyxin B:sodium oleate) charge ratio in the same donor mucus with display statistics indicating that the difference between the two formulations is significant according to a paired t-test. (C&D) Trajectories for each formulation. The data points in Figure 8B represent the individual values obtained from each slide. Error bars represent the standard deviation of *n* = 7 slides.

## Conclusion

The relationships among formulation parameters, internal liquid crystalline mesophase, and *in vitro* performance were systematically established for diblock copolymer stabilized nanocarriers co-encapsulating a hydrophilic peptide with one of four hydrophobic payloads. Across all formulations, increased ionic strength and decreased pH promoted decomplexation of the hydrophobic ion paired complexes responsible for driving liquid crystalline mesophase formation, resulting in small angle X-ray scattering profiles with lower peak intensities compared to the samples prepared in MilliQ water. These effects were most pronounced at the stoichiometric charge ratio, suggesting that the liquid crystalline mesophases formed at higher charge ratios are stabilized by the excess counterion and can therefore exhibit higher order structures even at lower pH. Independent of release medium, formulations containing vitamin E acetate, methyl oleate, or cholesterol produced liquid crystalline structures with larger characteristic d-spacings than those incorporating polycaprolactone, which exhibited d-spacings comparable to nanocarriers without a co-encapsulated payload. These structural differences correlated with polymyxin B release, with formulations exhibiting larger characteristic spacings also exhibited slower drug release. By comparing the release rate of polymyxin B from identical formulations in different media, new experimental support for the mechanisms that govern release of hydrophobic ion paired complexes from diblock copolymer stabilized nanocarrier cores was elucidated. These findings contextualize the results of bacterial efficacy studies wherein nanocarriers formulated at a 1:1 charge ratio achieved therapeutic performance comparable to free drug, while the slower release observed at a 1:4 charge ratio correlated with limited efficacy over the time scale of the assay.

Collectively, these findings establish a reproducible workflow for evaluating release kinetics in liquid crystalline nanocarrier systems and define structure–function relationships that inform the rational design of therapeutic carriers co-encapsulating hydrophilic and hydrophobic drugs. Future work should focus on applying analytical techniques such time-resolved small angle X-ray scattering of therapeutic nanocarriers over the course of drug release, small angle neutron scattering, and single particle analysis to validate the proposed mechanism for drug release attenuation in liquid crystalline nanocarriers. Other future studies should include performance *in vivo* against bacterial infection models in physiological environments such as the respiratory tract to delineate where the observed differences in release rate have the most biologically significant effects. Such studies would also benefit from the development of predictive tools to assess the favorability of interactions between ion paired complexes and co-encapsulated payloads to inform nanocarrier formulation design.

## Materials & Methods

### Materials

Polymyxin B (PMB) sulfate was purchased from Tokyo Chemical Industry America (Portland, OR, USA). Sodium oleate (NaOL) was purchased from Thermo Scientific (Waltham, MA, USA). NaOL was used as a hydrophobic counterion for ion pairing to polymyxin B for encapsulation. Polycaprolactone-b-poly(ethylene glycol) (PCL_5k_-b-PEG_5k_) was purchased from PolymerSource (Montreal, Quebec, Canada) and used the nanocarrier stabilizer. Polycaprolactone (2k) was purchased from PolymerSource (Montreal, Quebec, Canada). Vitamin E Acetate (97%, VEAc) was purchased from Thermo Scientific (Waltham, MA, USA). Methyl Oleate (MeOL) was purchased from Millipore Sigma (St. Louis, MO, USA). Cholesterol (CHL) was purchased from Thermo Scientific (Waltham, MA, USA). VEAc, CHL, MeOL, and PCL_2k_ were used as hydrophobic co-cores in this work. Tetrahydrofuran (THF) (HPLC grade, 99.9%) and methanol (MeOH, HPLC grade, 99.9% purity) for SAXS work was provided courtesy of the Australian Synchrotron, part of ANSTO. THF (HPLC grade), MeOH (HPLC grade), and acetonitrile (ACN) for formulating nanocarriers for performance experiments were purchased from Fisher Chemical (Loughborough, UK). Trifluoroacetic acid (TFA, LC/MS Grade) and formic acid (FA, LC/MS grade) were purchased from Fisher Chemical (Loughborough, UK) for preparation of mobile phases and extractions. Sodium hydroxide (97.0% Purity) and hydrochloric acid (12M) were purchased from Fisher Chemical (Loughborough, UK) and utilized to pH adjust release media. 10x phosphate buffered saline used for release tests was purchased from Fisher Chemical (Loughborough, UK). ((3-cholamidopropyl)dimethylammonio)-1-propanesulfonate (CHAPS) was purchased from Thermo Scientific (Waltham, MA, USA) and used as a hydrophobic sink in release testing. Water was MilliQ grade. HEPES free acid was purchased from Fisher Scientific (Waltham, MA, USA) and used to prepare the buffer dispersant media used when measuring nanocarrier zeta potential. Potassium chloride (99%, KCl) was purchased from Thermo Scientific (Waltham, MA, USA) and used to increase the conductivity of the dispersant media used when measuring nanocarrier zeta potential. Bacteria Pseudomonas aeruginosa 95 (ATCC 14211) was purchased from American Type Culture Collection (Manassas, VA, USA). Brain Heart Infusion (BHI) media used for bacterial culture was obtained from Becton Dickinson (Franklin Lakes, NJ, USA). Bacteriological agar was purchased from ThermoFisher Scientific (Rockford, IL, USA). Resazurin sodium was sourced from Medchem Express (Monmouth Junction, NJ, USA). Materials for mucus diffusion studies are provided where discussed.

### Nanocarrier formulation via Flash NanoPrecipitation

Flash NanoPrecipitation is an established, reproducible nanocarrier manufacturing technique that relies on rapid mixing to drive precipitation and self-assembly of nanocarrier components[20]. Prior to FNP, the hydrophobic co-core (vitamin E acetate, cholesterol, methyl oleate, or polycaprolactone at 10%, 25%, or 50% by mass of the total core), hydrophobic counterion (sodium oleate at 1:1, 1:2, or 1:4 polymyxin to counterion charge ratio), and stabilizer (PCL_5k_-b-PEG_5k_) were individually dissolved at 3x the desired feed stream concentration in an organic solvent, either MeOH or THF. The stock solutions were then combined in equal parts to yield a 1x feed stream. Polymyxin B was dissolved at 5 mg/mL using MilliQ (MQ) water. Aqueous and organic feed streams were combined in a confined impinging jets mixer (CIJ) by rapidly manually depressing Leur-slip syringes, as reported elsewhere[20]. The mixer effluent was collected in a 4 mL quench bath to dilute the concentration of organic solvent to 10% by volume and arrest nanocarrier growth. After manufacture, all LCNC suspensions were dialyzed overnight against MQ water to remove residual organic solvent. Following dialysis, LCNCs for synchrotron SAXS were concentrated 5x over Amicon filters (10k rcf, Millipore Sigma (St. Louis, MO, USA)) to improve data quality, in line with previous studies[22, 49]. Nanocarriers for mucus diffusivity assessment were frozen over 500 mM sucrose to prevent aggregation and degradation during transport. The size and surface charge of the LCNCs for *in vitro* work was determined pre– and post-dialysis using a Malvern Zetasizer Pro (Malvern Instruments, Great Malvern, England). Size was measured by diluting the nanocarrier suspension into water, and zeta potential was evaluated for LCNC suspensions diluted into 10mM pH 7.2 HEPES buffer supplemented with KCl to ensure sufficient conductivity (1-2 mS/cm). The HEPES buffer was titrated to the target pH using 1M NaOH prior to adding the final volume of water required to achieve the desired concentration.

The size of the LCNCs for synchrotron SAXS was measured for fresh, dialyzed, and concentrated samples using a Malvern Zetasizer Nano (Malvern Instruments, Great Malvern, England). After concentration, 30 uL of media (MQ, 10x PBS at pH 4.5 (nominal pH 4.54 [H+] = 0.0045 M), or 10x PBS at pH 7.4 (pH 7.49) was added to 270 uL of NCs. The NCs were exposed to each condition (MQ, PBS with nominal pH 4.5 (160 mM PBS, 0.0045M HCl), or PBS at pH 7.4) for 72 hr prior to measurement to allow for equilibration. Size data are provided in the supplemental information (SI) **Section 1**.

### Encapsulation efficiency

Encapsulation efficiency (EE) of LCNC formulations was calculated after dialysis via extraction and high-performance liquid chromatography (HPLC). For the extraction protocol, 300 µL of dialyzed LCNCs were added to 400 µL ACN with 0.1% formic acid. The resulting mixture was incubated at room temperature for 15 mins prior to filtration (Watman Puradisc 13 mm, Cytiva, Marlborough, MA) into HPLC vials. Controls of free PMB and PMB post FNP were performed to determine the efficiency of the protocol and are provided in the supplemental information (**SI, Section 4.2**). The EE of LCNCs was determined using equation 1:

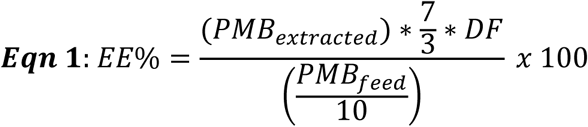

In equation 1, PMB_extracted_ is the concentration of PMB determined using the calibration curve and HPLC readout (mAU) in mg/mL, PMB_feed_ is the concentration of PMB in the feed stream in mg/mL, and DF is the dilution factor determined using the volume change over dialysis. For release testing, the value determined for the concentration of PMB post-dialysis (PMB_extracted_) was utilized as the initial concentration of PMB in each vessel. If the HPLC trace showed no peak or a peak below the lowest point on the standard curve, the area obtained for the lowest point on the standard curve was used for data analysis. All formulations that share the same co-core weight fraction and charge ratio also exhibit the same drug loading summarized in the table below.

**Table 1:** Summary of hydrophilic and hydrophobic API drug loadings (DL) by concentration (mg/mL API / mg/mL total formulation) for all formulations presented herein. The concentration of hydrophilic active in all formulations is the same (0.5 mg/mL) and the concentration of the hydrophobic co-core was adjusted based on total core composition such that the weight fraction of the hydrophobic co-core was equal to 10%, 25%, or 50% of the total core (PMB + NaOL).

| Charge ratio | Co-core weight fraction | Polymyxin B DL (% w/w) |
| --- | --- | --- |
| 1:1 | 0% | 32% |
| 1:1 | 10% | 29% |
| 1:1 | 25% | 26% |
| 1:1 | 50% | 19% |
| 1:4 | 0% | 15% |
| 1:4 | 10% | 14% |
| 1:4 | 25% | 12% |
| 1:4 | 50% | 8% |

### High performance liquid chromatography

All analytes of interest (PMB and VEAc) were quantified using an Agilent 1290 Infinity HPLC system with a diode array detector. PMB concentration was measured using a Waters XSelect HSS T3 Column, 100Å, 3.5 µm, 4.6 mm X 150 mm. An acetonitrile/water/TFA gradient method was employed to detect PMB at a total flow rate of 1 mL/min, column temperature of 27°C, and detection at 220 nm as reported previously[50]. To quantify VEAc, an isocratic method using a 99:1 v:v Methanol:Water mobile phase at a flow rate of 1.4 mL/min was adopted from Navale *et al.* to a Phenomenex Gemini 5 µm C18 110 Å, LC Column (150 x 4.6 mm)[51]. The detector wavelength was set to 292 nm and no column temperature control was utilized. All injection volumes were 20 µL. The standard curves utilized for experimental analyses are provided in the SI.

### Synchrotron small angle X-ray scattering data collection

SAXS data were collected using the automated sample handler at the Small and Wide Angle X-ray Scattering (SAXS/WAXS) beamline at the Australian Synchrotron, part of ANSTO[52]. Measurements were performed under ambient conditions (27°C) in the experimental hutch where the beamline was located. SAXS data were collected in the *q*-range 0.01-0.8 Å^-1^ using a maximum X-ray energy of 12 keV (wavelength = 1.0322 Å) and a detector distance of 2220.1 mm. The scattering vector, *q*, is equivalent to 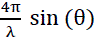 where θ represents the scattering angle[53–55]. Between formulations, blank measurements were taken with water to check for fouling of the capillary. The scattering pattern recorded by the Pilatus 2M detector was integrated into 1D plots of the intensity of the scattered X-rays I(*q*) vs *q* by the in-house program ScatterBrain. ScatterBrain X-Y data were exported into MATLAB for analysis and figure generation. Before export, the background (MQ water, PBS pH 7.4, or PBS nominal pH 4.5 (160 mM PBS, 0.0045M HCl)) was subtracted from the sample scattering patterns. Background subtracted SAXS patterns were imported to MATLAB for peak indexing via an in-house software. Identified Bragg peaks in the 2D plot of intensity versus *q*, representative of the reflection plane(s) within an LC phase, were related to one another to identify the internal structure of a given sample. A lamellar phase, for example, exhibits a √1:√4:√9:√16 ratio corresponding with *q*_1_:*q*_2_:*q*_3_:*q*_4[53–55]_. The d-spacing (d_spacing_) for each LC phase was calculated using a simplified version of Bragg’s law provided as **Equation 2**, where *q* is the location of the primary peak in Å^-1^ determined from the scattering profile[29, 53–55].

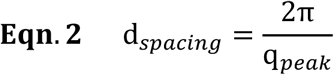

### *In vitro* release testing

Release media was prepared in bulk by diluting 10x PBS 9.9-fold, pH adjusting with HCl or NaOH, then adding additional volume such that the final solution contained 1x PBS titrated to the target pH. For media compositions containing a hydrophobic sink, CHAPS was added prior to pH adjusting such that the final concentration would be 10 mM. These pH values were selected based on two common pHs nanocarriers experience *in vivo*: the physiological pH of blood (pH 7.4) and the pH of lysosomes (pH 4.5)[56, 57]. CHAPS micelle formation at this concentration and ionic strength was verified using dynamic light scattering (**Figure S.9**). Release media was incubated overnight at 37°C to allow for temperature equilibration prior to the start of release testing. One hour prior to test initialization, clean beakers were filled with 350 mL of media and 100 kDa Float-A-Lyzer® devices (Repligen, Waltham, MA, USA) were inserted. The beakers were covered and incubated at 37°C again to allow for temperature equilibration. At t = 0 hr, 3.5 mL of the test sample was added to the Float-A-Lyzer® devices. At t = 1hr, 6hr, 12 hr, 24 hr, 72 hr, and 5d, aliquots were removed for analysis. 300 µL of the aliqout was removed for PMB extraction as described previously. The remaining volume (200 µL) was set aside (no co-core samples, PCL, and MeOL) or dried overnight under air (VEAc). After drying, 0.6 mL of organic solvent (THF) was added to the aliquot to solubilize the LCNC and extract the hydrophobic co-core. Samples were sonicated for 15 mins to increase the efficiency of the extraction prior to filtration.

### Efficacy against Pseudomonas aeruginosa

#### (i) Preparation of bacterial inoculum

The frozen bacterial stock of *P. aeruginosa* 95 (ATCC 14211) was streaked onto agar plates (1.5% w/v agar in BHI media) and was incubated overnight at 37°C to allow for colony formation. A single colony was isolated and inoculated into 100 mL BHI media followed by 16-hour incubation in an orbital shaker (100 rpm, 37°C). An aliquot of bacterial suspension (1 mL) was washed thrice with DPBS by centrifugation (2500 × *g,* 5 minutes, 4°C). The optical density (OD_600_) of bacterial suspension was measured and adjusted to 0.15 with BHI media, corresponding to 4 × 10^9^ CFU/mL.

#### (ii) Measurement of anti-bacterial efficacy using optical density and resazurin assay

The anti-bacterial efficacy was evaluated by measuring both optical density (OD_600_) and bacterial metabolic activity using a fluorescent dye, resazurin as an indicator[58]. Briefly, bacterial inoculum from step (i) was diluted to 5 × 10^5^ CFU/mL, and 100 mL of this suspension was incubated with 100 mL of the treatment in a 96-well non-treated plate (Corning, Durham, NC, USA). The free drug and formulations were diluted to achieve final PMB concentrations ranging from 0.25 – 5 mg/mL. Plates were incubated for 24 hours, at 37°C. The optical density (OD_600_) was measured before and after incubation using a Biotek Powerwave XS2 plate reader (Biotek Instruments, Inc., Winooski, VT, USA). To determine viable bacterial concentration (CFU/mL) after treatment, serial dilutions (10^2^ – 10^4^) were prepared and 50 mL of the respective dilution was streaked on the agar petri-plates followed by overnight incubation. Resazurin assay was performed to determine the minimum inhibitory concentration (MIC). After 24-hour incubation of treatments with the bacteria, 25 mL of resazurin dye (0.3 mg/mL) was added to each well. The plates were re-incubated at 37°C for 90 minutes, to allow the dye to interact with the bacteria. The color changes serve as an indicator for metabolic activity of the bacteria. Blue color indicates no bacterial growth, while pink indicates bacterial growth.

### Mucus diffusivity evaluation

Mucus collected from normal human bronchial epithelial (NHBE) cells grown at air-liquid interface (ALI) was provided courtesy of Dr. Margaret Scull at the University of Maryland. The mucus samples were frozen at –80°C until required experimentation. Prior to analysis, the mucus samples were thawed on ice along with the nanocarriers. Upon thawing, nanocarrier suspensions were diluted 1:10 in ultrapure water. For measurements, a microscope slide was prepared with a 25mm O-ring and sealed with vacuum grease. Each slide contained 24μL of thawed NHBE and 1μL of diluted NPs. The slide was then covered with a glass coverslip and left to equilibrate at 25°C for 30 minutes prior to imaging. A Zeiss 800 LSM microscope with a 63x water immersion lens and Zeiss Axiocam 702 camera were utilized to capture 10 second videos of the prepared microscope slide with a frame rate of 33.33 Hz. Multiple videos (≥5) were taken of each slide in distinct regions of the mucus. To extract exact time-dependent location data, the videos were run through a MATLAB script that calculates the time-averaged mean squared displacement (MSD) as a function of time scale, *τ*:

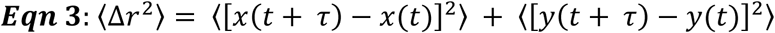

A separate MATLAB script was utilized to map nanocarrier trajectories in the mucus over each time period. Subsequent analysis and data visualization of MSD values was performed in Excel.

**Figure 9:**
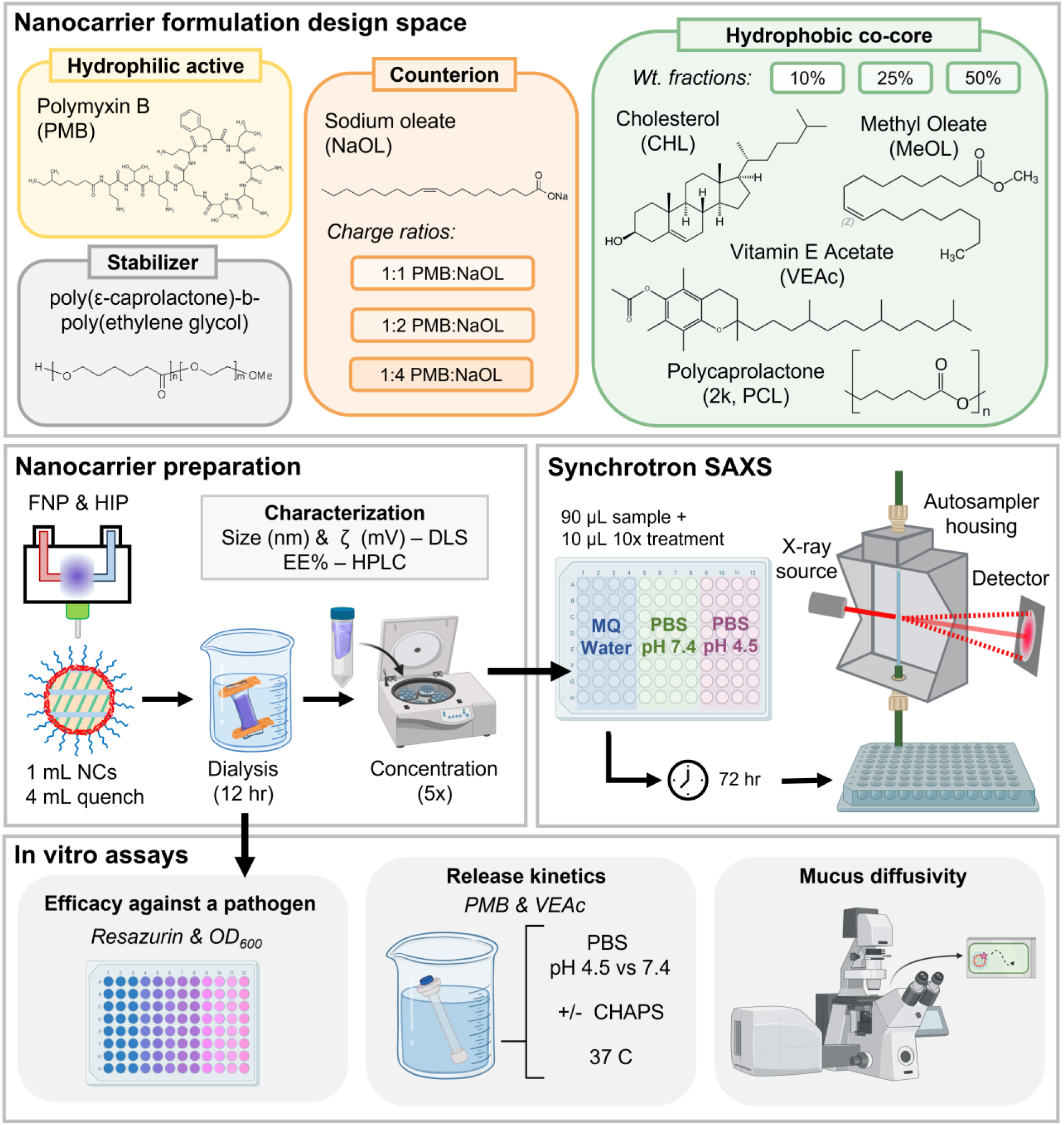
(Top) Chemical structures and formulation parameters covered in the nanocarrier design space. (Middle) Formulation preparation and characterization workflows in connection to (Bottom) *in vitro* assays. Portions of this figure were created with BioRender.com.

## Acknowledgements

The SAXS experiments for this work were conducted on the SAXS/WAXS beamline of the Australian Synchrotron, part of ANSTO. Beamtime award numbers were AS232/SAXS/19900 and AS232/SAXS/20059. Qi T Zhou and Shruti S. Sawant were supported by the National Institutes of Health under Award Numbers of R01AI146160 and R01HL167828. The content is solely the responsibility of the authors and does not necessarily represent the official views of the National Institutes of Health

This work was supported by a research grant by Serán Bioscience, LLC (to KDR). The authors thank Mark Kastantin, Ryan Mowry, and Tanner Corrado for insightful discussions around experimental design, data analysis, and manuscript construction. The authors also thank Anas Aljabbari, Elizabeth Edwards, and Emily Aicher for valuable conversations regarding protocol development and data presentation.

## Author contributions: CRediT

Sophia R. Dasaro (Methodology, Investigation, Formal analysis, writing – original draft)

Shruti S. Sawant (Methodology, Investigation, Formal analysis, writing – review & editing)

Alexa Stern (Methodology, Investigation, Formal analysis, writing – review & editing)

Lucas D. Johnson (Methodology, Investigation, Writing – review & editing)

Caleb F. Fretz (Methodology, Investigation, Writing – review & editing)

Malinda Salim (Methodology, Investigation, Writing – review & editing)

Nigel Kirby (Methodology, Investigation, Supervision

Ben J. Boyd (Resources, Writing – review & editing)

Brian K. Wilson (Conceptualization, Writing – review & editing)

Gregg Duncan (Resources, Writing – review & editing)

Qi Zhou (Resources, Writing – review & editing)

Kurt Ristroph (Conceptualization, Writing – review & editing, Funding acquisition)

## Data availability

The authors can make raw data available upon request.

