## Supporting information for "Liquid crystalline mesophase spacing as a quantitative predictor of release kinetics for co-loaded hydrophilic and hydrophobic payloads"

5: Australian Synchrotron, ANSTO, 800 Blackburn Road, Clayton, VIC 3168, Australia

6: Department of Pharmacy, University of Copenhagen, Universitetsparken 2, 2100 Copenhagen, Denmark

7: Bindley Bioscience Center, Purdue University, West Lafayette, Indiana, United States.

8: Davidson School of Chemical Engineering (by courtesy), Purdue University, 480 Stadium Mall Drive, West Lafayette, IN 47907, United States.

#### S.1 Size data of nanocarriers prepared for SAXS experiments

Nanocarriers for SAXS analysis were prepared at n = 1 with the size measured thought all processing stages as presented in **Table S.1** on the following page.

**Table S.1:** Size data for the nanocarrier formulations presented in the main text after FNP (Fresh), after dialysis (dialyzed), and post concentration (centrifugated). FNP stream compositions are provided in the white columns.

| June, 2023 | | Organic phase (mg/mL) | | | AQ (mg/mL) | Fresh | | Dialyzed | | Centrifuged | |
|---|---|---|---|---|---|---|---|---|---|---|---|
| **FNP#** | **Sample** | **NaOL** | **CC** | **PCL-PEG** | **PMB** | ***Size [nm]*** | ***PDI*** | ***Size [nm]*** | ***PDI*** | ***Size [nm]*** | ***PDI*** |
| 1 | 1:1, 0% cc | 5.85 | 0 | 5 | 5 | 150.0 | 0.07 | 202.3 | 0.13 | 194.5 | 0.09 |
| 2 | 1:1, 10% VitE | 5.85 | 1.21 | 5 | 5 | 95.1 | 0.10 | 137.9 | 0.22 | 148.1 | 0.14 |
| 3 | 1:1, 25% VitE | 5.85 | 2.62 | 5 | 5 | 75.8 | 0.15 | 125.3 | 0.23 | 139.0 | 0.20 |
| 4 | 1:1, 50% VitE | 5.85 | 10.9 | 5 | 5 | 57.5 | 0.12 | 69.1 | 0.27 | 86.7 | 0.13 |
| 5 | 1:1, 10% CHL | 5.85 | 1.21 | 5 | 5 | 97.0 | 0.15 | 122.7 | 0.18 | 147.1 | 0.13 |
| 6 | 1:1, 25% CHL | 5.85 | 2.62 | 5 | 5 | 94.3 | 0.09 | 134.1 | 0.09 | 162.0 | 0.14 |
| 8 | 1:1, 10% MeOL | 5.85 | 1.21 | 5 | 5 | 100.5 | 0.12 | 175.2 | 0.11 | 185.5 | 0.14 |
| 9 | 1:1, 25% MeOL | 5.85 | 2.62 | 5 | 5 | 81.9 | 0.08 | 199.3 | 0.08 | 210.9 | 0.12 |
| 10 | 1:1, 50% MeOL | 5.85 | 10.9 | 5 | 5 | 107.4 | 0.10 | 168.5 | 0.06 | 177.6 | 0.08 |
| 11 | 1:1, 10% PCL | 5.85 | 1.21 | 5 | 5 | 109.0 | 0.09 | 186.1 | 0.19 | 227.4 | 0.33 |
| 12 | 1:1, 25% PCL | 5.85 | 2.62 | 5 | 5 | 117.3 | 0.11 | 205.1 | 0.25 | 208.7 | 0.32 |
| 14 | 1:2, 0% cc | 11.7 | 0 | 5 | 5 | 129.4 | 0.15 | 418.2 | 0.15 | 297.7 | 0.26 |
| 15 | 1:2, 10% VitE | 11.7 | 1.85 | 5 | 5 | 89.1 | 0.14 | 208.5 | 0.10 | 215.6 | 0.14 |
| 16 | 1:2, 25% VitE | 11.7 | 5.85 | 5 | 5 | 76.5 | 0.12 | 127.2 | 0.14 | 174.8 | 0.07 |
| 17 | 1:2, 50% VitE | 11.7 | 16.7 | 5 | 5 | 75.5 | 0.13 | 90.9 | 0.23 | 129.5 | 0.14 |
| 18 | 1:2, 10% CHL | 11.7 | 1.85 | 5 | 5 | 97.9 | 0.13 | 212.6 | 0.09 | 256.6 | 0.23 |
| 19 | 1:2, 25% CHL | 11.7 | 5.85 | 5 | 5 | 101.6 | 0.10 | 244.6 | 0.11 | 295.7 | 0.40 |
| 21 | 1:2, 10% MeOL | 11.7 | 1.85 | 5 | 5 | 95.8 | 0.14 | 168.3 | 0.06 | 248.5 | 0.15 |
| 22 | 1:2, 25% MeOL | 11.7 | 5.85 | 5 | 5 | 85.8 | 0.08 | 188.7 | 0.10 | 206.5 | 0.09 |
| 23 | 1:2, 50% MeOL | 11.7 | 16.7 | 5 | 5 | 73.5 | 0.09 | 155.9 | 0.09 | 200.6 | 0.11 |

| | | | | | | | | | | | |
|---|---|---|---|---|---|---|---|---|---|---|---|
| 24 | 1:2, 10% PCL | 11.7 | 1.85 | 5 | 5 | 116.6 | 0.17 | 247.1 | 0.19 | 535.4 | 0.39 |
| 25 | 1:2, 25% PCL | 11.7 | 5.85 | 5 | 5 | 114.3 | 0.15 | 468.7 | 0.39 | 513.8 | 0.48 |
| 27 | 1:4, 0% cc | 23.4 | 0 | 5 | 5 | 126.4 | 0.16 | 149.3 | 0.15 | 141.6 | 0.13 |
| 28 | 1:4, 10%VE | 23.4 | 3.16 | 5 | 5 | 101.9 | 0.12 | 116.3 | 0.17 | 123.4 | 0.19 |
| 29 | 1:4, 25% VE | 23.4 | 9.49 | 5 | 5 | 81.9 | 0.10 | 94.3 | 0.22 | 110.1 | 0.22 |
| 30 | 1:4, 50% VE | 23.4 | 28.5 | 5 | 5 | 86.5 | 0.07 | 97.0 | 0.10 | 94.3 | 0.11 |
| 31 | 1:4, 10% CHL | 23.4 | 3.16 | 5 | 5 | 132.0 | 0.09 | 142.4 | 0.12 | 141.5 | 0.13 |
| 34^ | 1:4, 25% MeOL | 23.4 | 9.49 | 5 | 5 | 84.7 | 0.10 | 102.9 | 0.19 | 127.3 | 0.25 |
| 36 | 1:4, 50% MeOL | 23.4 | 28.5 | 5 | 5 | 85.4 | 0.09 | 116.5 | 0.13 | 114.8 | 0.10 |
| 37 | 1:4, 10% PCL | 23.4 | 3.16 | 5 | 5 | 134.8 | 0.11 | 147.8 | 0.14 | 136.5 | 0.14 |

1

### S.2 Supplemental SAXS data, profiles & tabulated peak indexing

*S2.1 SAXS data grouped by single formulation instead of charge ratio and condition*

Figures S.1-S.3 are included here to supplement the main text. Each panel contains the consists of one formulation after exposure to each of the three reported conditions.

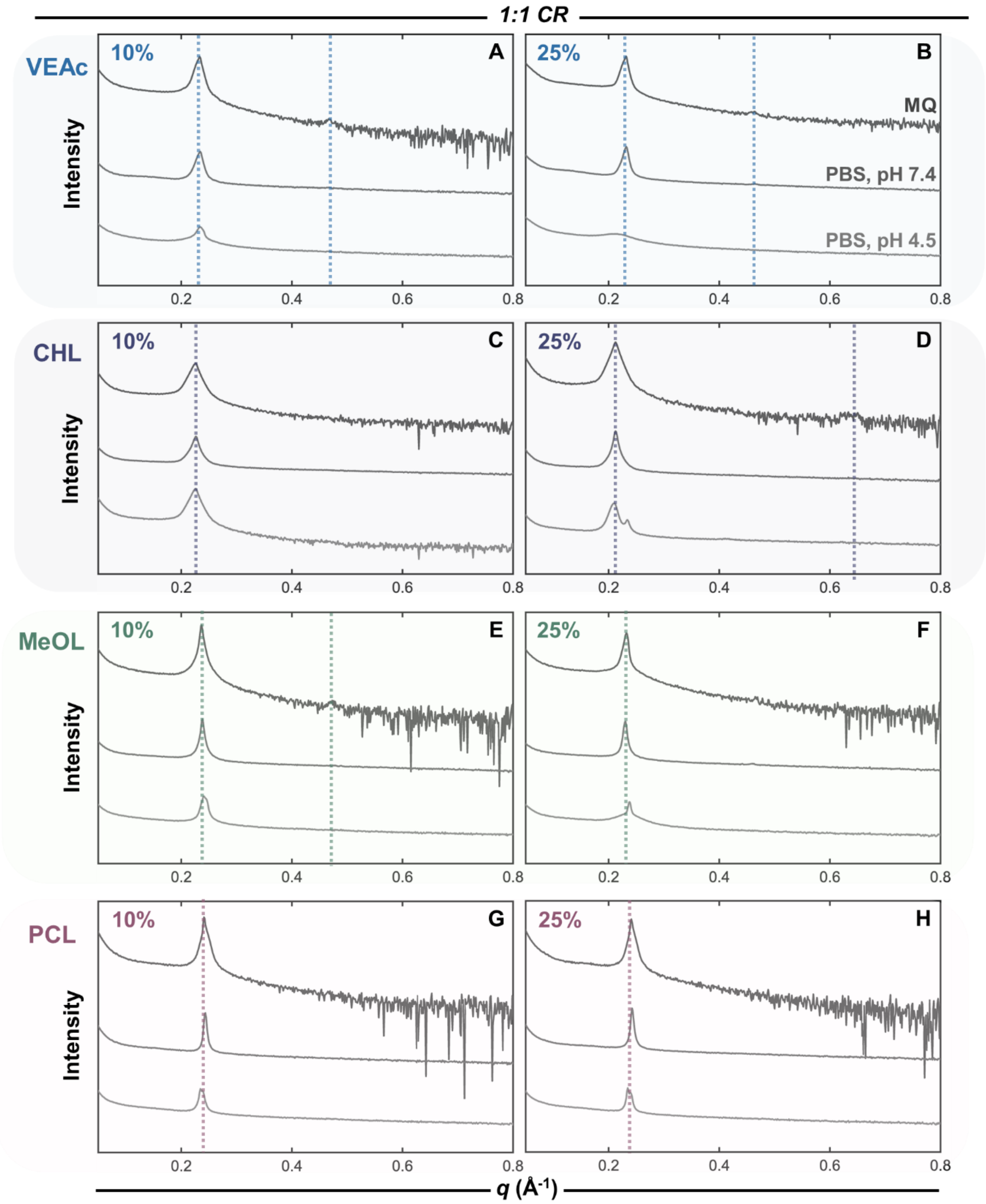

**Figure S.1**: PMB:OL LCNCs at a 1:1 charge ratio grouped by hydrophobic co-core.

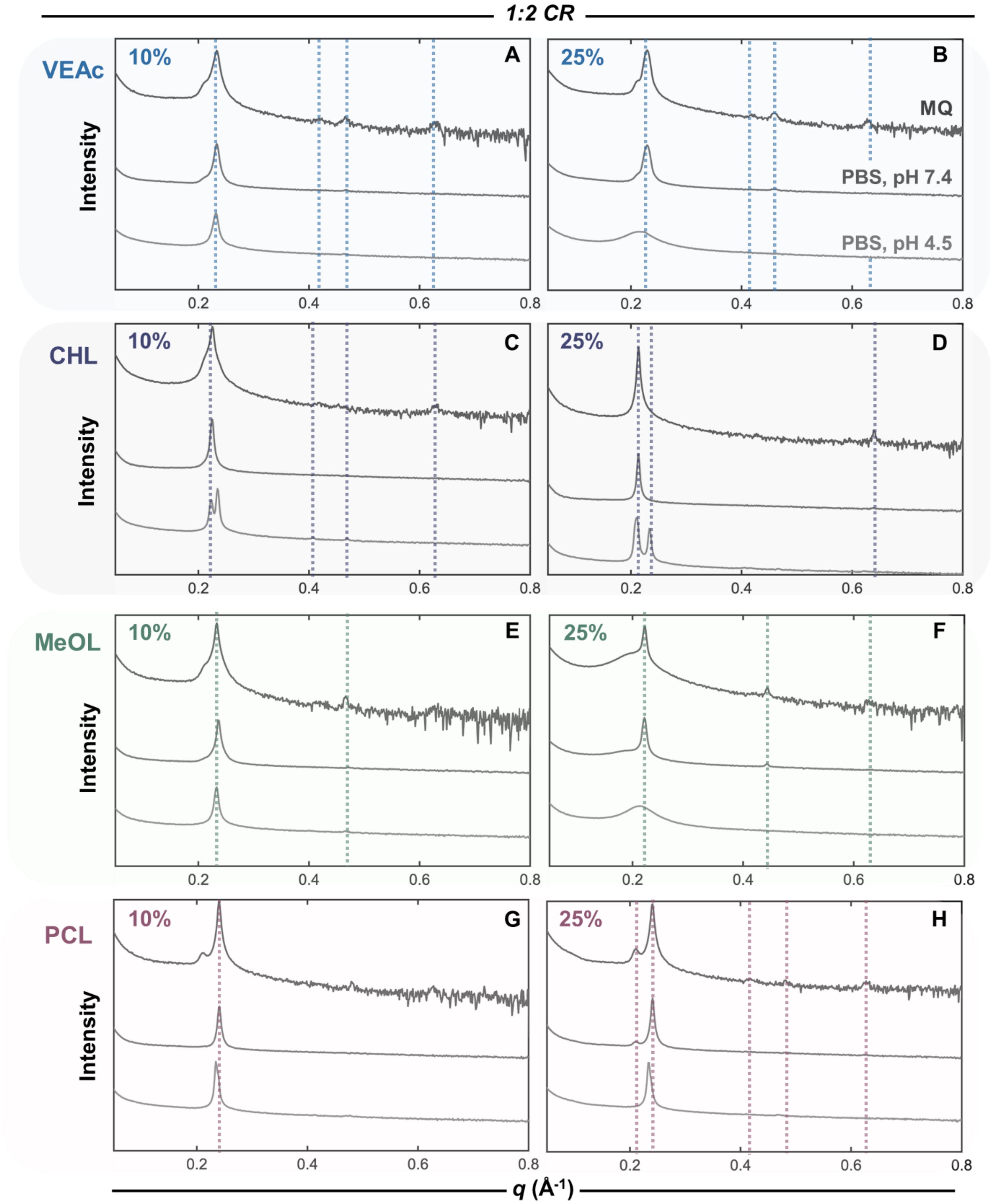

**Figure S.2**: PMB:OL LCNCs at a 1:1 charge ratio grouped by hydrophobic co-core

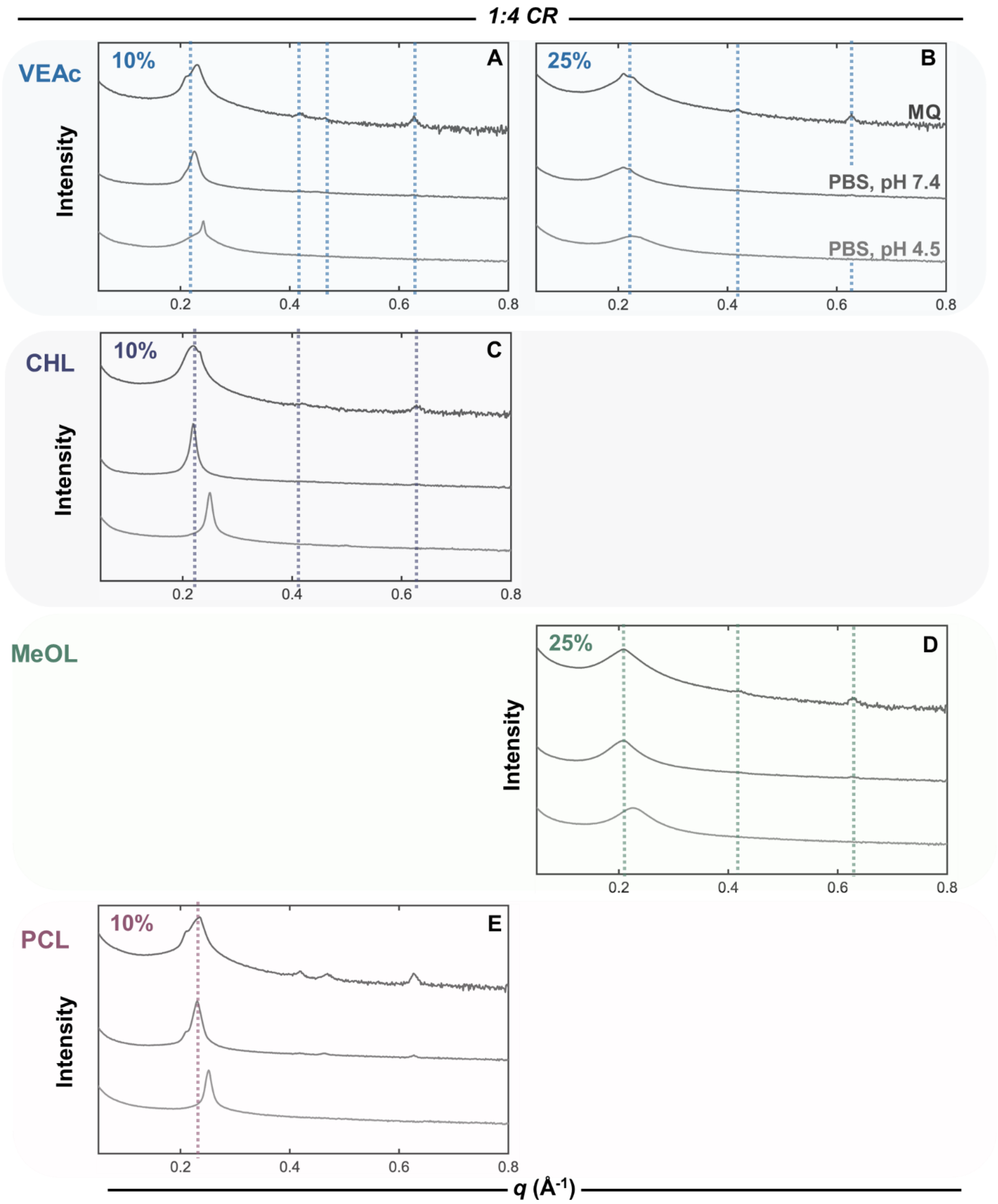

1 **Figure S.3**: PMB:OL nanocarriers at a 1:4 charge ratio grouped by hydrophobic co-core.

2

3

*S2.2 Grouped analysis for 25% wt. co-core*

**Figure S.4** presented below is the style equivalent of Figure 1 in the main body but for LCNCs formulated with a 25% wt. co-core.

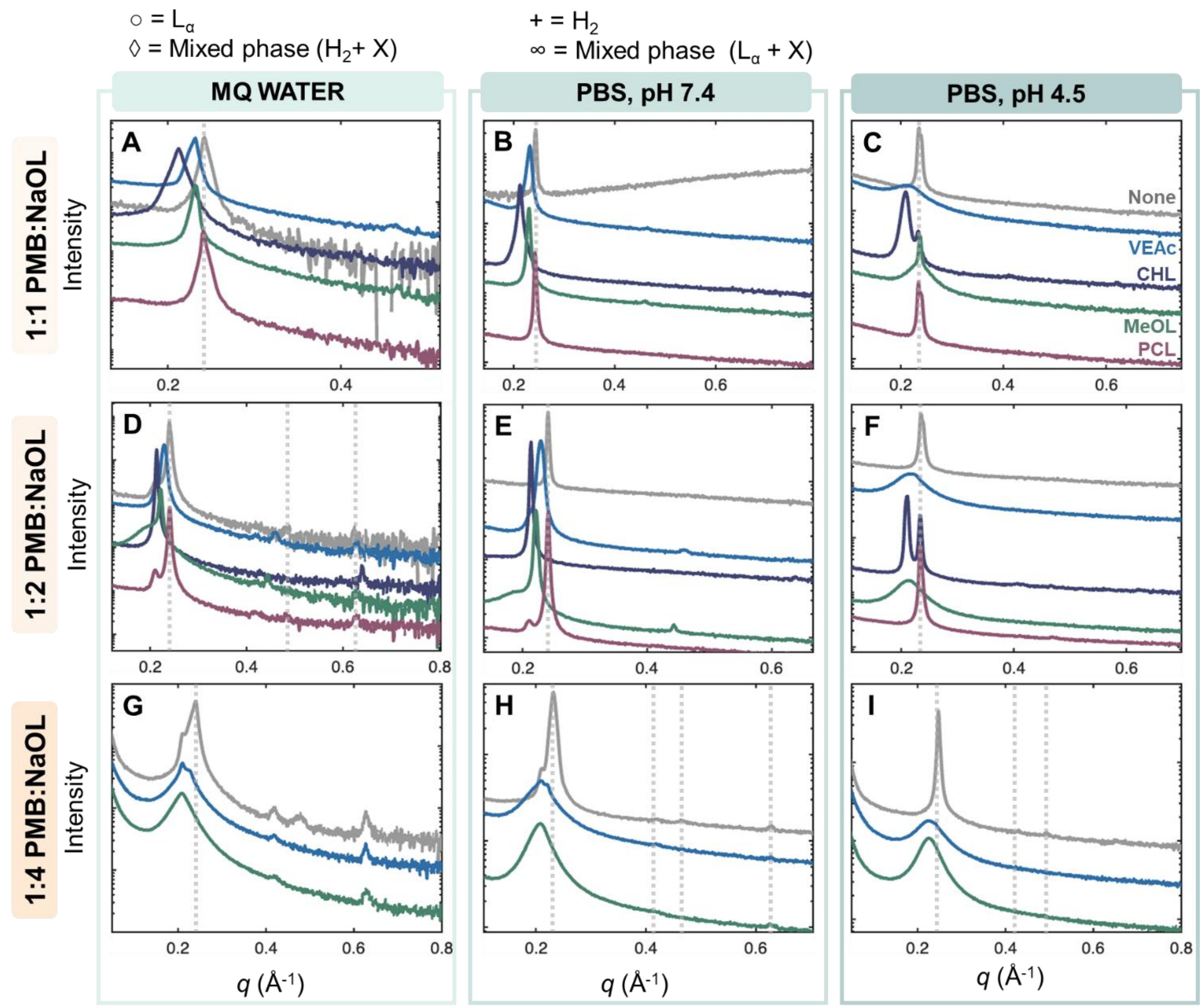

**Figure S.4:** SAXS profiles of liquid crystalline nanocarriers containing 0% or 10% of each hydrophobic co-core at a 1:1 (A-C), 1:2 (D-F), or 1:4 (G-I) charge ratio at all three conditions. Co-core type is color coded. Absence of a co-core trace denotes failure to generate process-stable NCs. Where identifiable, LC phases are denoted by symbols: ◊ = Mixed phase where one identified phase is inverse hexagonal ($H_2$), + = Inverse hexagonal, ○ = Lamellar ($L_\alpha$), and ∞ = Mixed phase where one identified phase is lamellar.

*S2.3 Tabulated summary of identified liquid crystalline phases*

**Table S.2**: Summary of the formulations and identified internal structures, represented by symbols in the main text. Structure was determined from the peak indices presented in **Tables S.3-6.**

| FNP# | Sample | MQ | PBS pH 7.4 | PBS pH 4.5 |
|---|---|---|---|---|
| 1 | 1:1, 0% cc | Undetermined | Undetermined | Mixed phase |
| 2 | 1:1, 10% VitE | Lamellar | Undetermined | Undetermined |
| 3 | 1:1, 25% VitE | Lamellar | Lamellar | Undetermined |
| 4 | 1:1, 50% VitE | Undetermined | Undetermined | Undetermined |
| 5 | 1:1, 10% CHL | Undetermined | Undetermined | Lamellar |
| 6 | 1:1, 25% CHL | Lamellar | Undetermined | Mixed phase (Inv. Hex + X) |
| 8 | 1:1, 10% MeOL | Lamellar | Undetermined | Mixed phase |
| 9 | 1:1, 25% MeOL | Lamellar | Lamellar | Undetermined |
| 10 | 1:1, 50% MeOL | Undetermined | Undetermined | Undetermined |
| 11 | 1:1, 10% PCL | Undetermined | Undetermined | Mixed phase |
| 12 | 1:1, 25% PCL | Undetermined | Undetermined | Mixed phase |
| 14 | 1:2, 0% cc | Mixed phase (Lamellar + X) | Undetermined | Undetermined |
| 15 | 1:2, 10% VitE | Mixed phase (Lamellar + X) | Lamellar | Lamellar |
| 16 | 1:2, 25% VitE | Mixed phase (Lamellar + X) | Lamellar | Undetermined |
| 17 | 1:2, 50% VitE | Mixed phase (Lamellar + X) | Lamellar | Undetermined |
| 18 | 1:2, 10% CHL | Mixed phase (Lamellar + X) | Undetermiend | Mixed phase (Inv. Hex + X) |
| 19 | 1:2, 25% CHL | Lamellar | Lamellar | Mixed phase (Inv. Hex + X) |
| 21 | 1:2, 10% MeOL | Mixed phase (Lamellar + X) | Lamellar | Mixed phase (Lamellar + X) |
| 22 | 1:2, 25% MeOL | Lamellar | Mixed phase (Lamellar + X) | Undetermined |
| 23 | 1:2, 50% MeOL | Lamellar | Lamellar | Undetermined |
| 24 | 1:2, 10% PCL | Mixed phase (Inv. Hex + X) | Undetermined | Lamellar |
| 25 | 1:2, 25% PCL | Mixed phase (Inv. Hex + X) | Undetermined | Lamellar |
| 27 | 1:4, 0% cc | Mixed phase (Inv. Hex + X) | Mixed phase (Lamellar + X) | Inverse hexagonal |
| 28 | 1:4, 10%VE | Mixed phase (Inv. Hex + X) | Mixed phase (Lamellar + X) | Undetermiend |
| 29 | 1:4, 25% VE | Mixed phase (Lamellar + X) | Mixed phase (Lamellar + X) | Undetermined |
| 30 | 1:4, 50% VE | Mixed phase | Undetermined | Undetermined |
| 31 | 1:4, 10% CHL | Mixed phase (Lamellar + X) | Undetermined | Lamellar |
| 34^ | 1:4, 10% meol | Mixed phase (Lamellar + X) | Lamellar | Undetermined |
| 36 | 1:4, 50% meol | Lamellar | Undetermined, Broad peak | Undetermined |
| 37 | 1:4, 10% PCL | Mixed phase (Inv. Hex + X) | Mixed phase (Lamellar + X) | Undetermined |

*S2.4 Peak indexing summary*

**Table S.3**: Summary of formulations, primary peak locations, and identifiable LC phases for samples in MQ water after 72 hr. ^ denotes NC formulations hypothesized to have crashed out prior to analysis (no long-term stability) but were resuspended and ran on the autoloader to complete the screen. For mixed phase samples where a dominant LC phase was identifiable, the phase is reported following: Main + X represents the unidentified phase due to peak overlap. Indexing was done to two start points and compared to one another in an effort to deconvolute samples where mixed LC phases were present.

| FNP# | Sample | Peak Locations | | | | | Peak Ratios (indexed to first) | | | | | Peak Ratios (indexed to largest) | | | | LC Phase |
|---|---|---|---|---|---|---|---|---|---|---|---|---|---|---|---|---|
| 1 | 1:1, 0% cc | 0.241 | | | | | 1.000 | | | | | | | | | Undetermined |
| 2 | 1:1, 10% VitE | 0.233 | 0.466 | | | | 1.000 | 2.000 | | | | | | | | Lamellar |
| 3 | 1:1, 25% VitE | 0.231 | 0.459 | | | | 1.000 | 1.987 | | | | | | | | Lamellar |
| 4 | 1:1, 50% VitE | 0.229 | | | | | 1.000 | | | | | | | | | Undetermined |
| 5 | 1:1, 10% CHL | 0.224 | | | | | 1.000 | | | | | | | | | Undetermined |
| 6 | 1:1, 25% CHL | 0.212 | 0.633 | | | | 1.000 | 2.986 | | | | | | | | Lamellar |
| 8 | 1:1, 10% MeOL | 0.237 | 0.475 | | | | 1.000 | 2.004 | | | | | | | | Lamellar |
| 9 | 1:1, 25% MeOL | 0.232 | 0.469 | | | | 1.000 | 2.022 | | | | | | | | Lamellar |
| 10 | 1:1, 50% MeOL | 0.185 | | | | | 1.000 | | | | | | | | | Undetermined |
| 11 | 1:1, 10% PCL | 0.241 | | | | | 1.000 | | | | | | | | | Undetermined |
| 12 | 1:1, 25% PCL | 0.242 | | | | | 1.000 | | | | | | | | | Undetermined |
| 14 | 1:2, 0% cc | 0.209 | 0.239 | 0.482 | 0.621 | | 1.000 | 1.144 | 2.306 | 2.971 | | 1.000 | 2.017 | 2.598 | | Mixed phase (Lamellar + X) |
| 15 | 1:2, 10% VitE | 0.209 | 0.233 | 0.424 | 0.466 | 0.627 | 1.000 | 1.115 | 2.029 | 2.230 | 3.000 | 1.000 | 1.820 | 2.000 | 2.691 | Mixed phase (Lamellar + X) |
| 16 | 1:2, 25% VitE | 0.209 | 0.228 | 0.416 | 0.462 | 0.627 | 1.000 | 1.091 | 1.990 | 2.211 | 3.000 | 1.000 | 1.825 | 2.026 | 2.750 | Mixed phase (Lamellar + X) |
| 17 | 1:2, 50% VitE | 0.209 | 0.228 | 0.457 | 0.623 | | 1.000 | 1.091 | 2.187 | 2.981 | | 1.000 | 2.004 | 2.732 | | Mixed phase (Lamellar + X) |
| 18 | 1:2, 10% CHL | 0.211 | 0.226 | 0.416 | 0.451 | 0.626 | 1.000 | 1.841 | 1.996 | 2.770 | | 1.000 | 1.841 | 1.996 | 2.770 | Mixed phase (Lamellar + X) |
| 19 | 1:2, 25% CHL | 0.214 | 0.427 | 0.639 | | | 1.000 | 1.995 | 2.986 | | | 1.000 | 1.496 | | | Lamellar |
| 21 | 1:2, 10% MeOL | 0.209 | 0.233 | 0.417 | 0.466 | 0.633 | 1.000 | 1.115 | | | | 1.000 | 1.790 | 2.000 | 2.717 | Mixed phase (Lamellar + X) |
| 22 | 1:2, 25% MeOL | 0.222 | 0.444 | | | | 1.000 | 2.000 | | | | | | | | Lamellar |
| 23 | 1:2, 50% MeOL | 0.219 | 0.437 | | | | 1.000 | 1.995 | | | | | | | | Lamellar |
| 24 | 1:2, 10% PCL | 0.209 | 0.239 | 0.422 | 0.479 | 0.627 | 1.000 | 1.144 | 2.019 | 2.292 | 3.000 | 1.000 | 1.766 | 2.004 | 2.623 | Mixed phase (Inv. Hex + X) |
| 25 | 1:2, 25% PCL | 0.208 | 0.241 | 0.420 | 0.481 | 0.627 | 1.000 | 1.159 | 2.019 | 2.313 | 3.014 | 1.000 | 1.743 | 1.996 | 2.602 | Mixed phase (Inv. Hex + X) |
| 27 | 1:4, 0% cc | 0.210 | 0.241 | 0.420 | 0.482 | 0.633 | 1.000 | 1.148 | 2.000 | 2.295 | 3.014 | 1.000 | 1.743 | 2.000 | 2.627 | Mixed phase (Inv. Hex + X) |
| 28 | 1:4, 10%VE | 0.208 | 0.240 | 0.417 | 0.474 | 0.628 | 1.000 | 1.154 | 2.005 | 2.279 | 3.019 | 1.000 | 1.738 | 1.975 | 2.617 | Mixed phase (Inv. Hex + X) |
| 29 | 1:4, 25% VE | 0.208 | 0.228 | 0.418 | 0.466 | 0.627 | 1.000 | 1.096 | 2.010 | 2.240 | | 1.000 | 1.833 | 2.044 | 2.750 | Mixed phase (Lamellar + X) |
| 30 | 1:4, 50% VE | 0.209 | 0.228 | 0.419 | 0.627 | | 1.000 | 1.091 | 2.005 | | | 1.000 | 1.838 | 2.750 | | Mixed phase |
| 31 | 1:4, 10% CHL | 0.209 | 0.418 | 0.627 | | | 1.000 | 2.000 | 3.000 | | | | | | | Mixed phase (Lamellar + X) |
| 32^ | 1:4, 25% CHL | 0.218 | 0.232 | 0.418 | 0.462 | 0.627 | 1.000 | 1.064 | 1.917 | 2.119 | 2.876 | 1.000 | 1.802 | 1.991 | 2.703 | Mixed phase |
| 33^ | 1:4, 50% CHL | 0.208 | 0.410 | 0.612 | | | 1.000 | 1.971 | 2.942 | | | 1.000 | 1.493 | | | Lamellar |
| 34^ | 1:4, 10% meol | 0.208 | 0.237 | 0.418 | 0.628 | | 1.000 | 1.139 | 2.010 | 3.019 | | 1.000 | 1.764 | 2.650 | | Mixed phase (Lamellar + X) |
| 35 | 1:4, 25% meol | 0.207 | 0.423 | 0.628 | | | 1.000 | 2.043 | 3.034 | | | 1.000 | 1.485 | | | Lamellar |
| 36 | 1:4, 50% meol | 0.204 | 0.421 | 0.627 | | | 1.000 | 2.065 | 3.074 | | | 1.000 | 1.489 | | | Lamellar |
| 37 | 1:4, 10% PCL | 0.210 | 0.235 | 0.418 | 0.427 | 0.627 | 1.000 | 1.119 | 1.990 | 2.033 | 2.986 | 1.000 | 1.779 | 1.817 | 2.668 | Mixed phase (Inv. Hex + X) |
| 38^ | 1:4, 25% PCl | 0.209 | 0.248 | 0.424 | 0.470 | 0.627 | 1.000 | 1.187 | 2.029 | 2.249 | 3.000 | 1.000 | 1.710 | 1.895 | 2.528 | Mixed phase (Inv. Hex + X) |
| 39^ | 1:4, 50% PCL | 0.208 | 0.237 | 0.420 | 0.477 | 0.627 | 1.000 | 1.139 | 2.019 | 2.293 | 3.014 | 1.000 | 1.772 | 2.013 | 2.646 | Mixed phase (Inv. Hex + X) |

**Table S.4**: Summary of formulations, primary peak locations, and identifiable LC phases for samples in PBS at pH 7.4 after 72 hr. Nomenclature is as previously described.

| FNP# | Sample | Peak Locations | | | | | Peak Ratios (indexed to first) | | | | | Peak Ratios (indexed to largest) | | | | LC Phase |
|---|---|---|---|---|---|---|---|---|---|---|---|---|---|---|---|---|
| 1 | 1:1, 0% | 0.244 | | | | | 1.000 | | | | | | | | | Undetermined |
| 2 | 1:1, 10% VE | 0.234 | | | | | 1.000 | | | | | | | | | Undetermined |
| 3 | 1:1, 25% VE | 0.232 | 0.465 | | | | 1.000 | 2.004 | | | | | | | | Lamellar |
| 4 | 1:1, 50% VE | 0.231 | | | | | 1.000 | | | | | | | | | Undetermined |
| 5 | 1:1, 10% Chol | 0.226 | | | | | 1.000 | | | | | | | | | Undetermined |
| 6 | 1:1, 25% Chol | 0.212 | | | | | 1.000 | | | | | | | | | Undetermined |
| 8 | 1:1, 10% MeOL | 0.238 | | | | | 1.000 | | | | | | | | | Undetermined |
| 9 | 1:1, 25% MeOL | 0.230 | 0.460 | | | | 1.000 | 2.000 | | | | | | | | Lamellar |
| 10 | 1:1, 50% MeOL | 0.190 | | | | | 1.000 | | | | | | | | | Undetermined |
| 11 | 1:1, 10% PCL | 0.243 | | | | | 1.000 | | | | | | | | | Undetermined |
| 12 | 1:1, 25% PCL | 0.242 | | | | | 1.000 | | | | | | | | | Undetermined |
| 14 | 1:2, 0% | 0.241 | | | | | 1.000 | | | | | | | | | Undetermined |
| 15 | 1:2, 10% VE | 0.233 | 0.468 | | | | 1.000 | 2.009 | | | | 1.000 | | | | Lamellar |
| 16 | 1:2, 25% VE | 0.229 | 0.460 | | | | 1.000 | 2.009 | | | | 1.000 | | | | Lamellar |
| 17 | 1:2, 50% VE | 0.228 | 0.456 | | | | 1.000 | 2.000 | | | | 1.000 | | | | Lamellar |
| 18 | 1:2, 10% Chol | 0.225 | | | | | 1.000 | | | | | | | | | Undetermiend |
| 19 | 1:2, 25% Chol | 0.214 | 0.639 | | | | 1.000 | 2.986 | | | | 1.000 | | | | Lamellar |
| 21 | 1:2, 10% MeOL | 0.236 | 0.474 | | | | 1.000 | 2.008 | | | | 1.000 | | | | Lamellar |
| 22 | 1:2, 25% MeOL | 0.184 | 0.222 | 0.442 | | | 1.000 | 1.207 | 2.402 | | | 1.000 | 1.991 | | | Mixed phase (Lamellar + X) |
| 23 | 1:2, 50% MeOL | 0.219 | 0.437 | | | | 1.000 | 1.995 | | | | | | | | Lamellar |
| 24 | 1:2, 10% PCL | 0.241 | | | | | 1.000 | | | | | | | | | Undetermined |
| 25 | 1:2, 25% PCL | 0.211 | 0.241 | | | | 1.000 | 1.142 | | | | 1.000 | | | | Undetermined |
| 27 | 1:4, 0% | 0.210 | 0.232 | 0.421 | 0.464 | 0.628 | 1.000 | 1.105 | 2.005 | 2.210 | 2.990 | 1.000 | 1.815 | 2.000 | 2.707 | Mixed phase (Lamellar + X) |
| 28 | 1:4, 10% VE | 0.225 | 0.418 | 0.451 | 0.627 | | 1.000 | 1.858 | 2.004 | 2.787 | | 1.000 | 1.079 | 1.500 | | Mixed phase (Lamellar + X) |
| 29 | 1:4, 25% VE | 0.210 | 0.222 | 0.418 | 0.627 | | 1.000 | 1.057 | 1.990 | 2.986 | | 1.000 | 1.883 | 2.824 | | Mixed phase (Lamellar + X) |
| 30 | 1:4, 50% VE | 0.207 | 0.228 | | | | 1.000 | 1.101 | | | | 1.000 | | | | Undetermined |
| 31 | 1:4, 10% Chol | 0.219 | | | | | 1.000 | | | | | | | | | Undetermined |
| 35 | 1:4, 25% MeOL | 0.208 | 0.628 | | | | 1.000 | 3.019 | | | | 1.000 | | | | Lamellar |
| 36 | 1:4, 50% MeOL | 0.180 | 0.211 | | | | 1.000 | 1.172 | | | | 1.000 | | | | Undetermined, Broad peak |
| 37 | 1:4, 10% PCL | 0.210 | 0.231 | 0.420 | 0.465 | 0.627 | 1.000 | 1.100 | 2.000 | 2.214 | | 1.000 | 1.818 | 2.013 | | Mixed phase (Lamellar + X) |

**Table S.5**: Summary of formulations, primary peak locations, and identifiable LC phases for samples in PBS at pH 4.5 after 72 hr. Nomenclature is as previously described.

| FNP# | Sample | Peak Locations | | | | | Peak Ratios (indexed to first) | | | | | Peak Ratios (indexed to largest) | | | | LC Phase |
|---|---|---|---|---|---|---|---|---|---|---|---|---|---|---|---|---|
| 1 | 1:1, 0% | 0.234 | 0.240 | | | | 1.000 | 1.026 | | | | 1.000 | | | | Mixed phase |
| 2 | 1:1, 10% VE | 0.235 | | | | | 1.000 | | | | | | | | | Undetermined |
| 3 | 1:1, 25% VE | 0.214 | | | | | 1.000 | | | | | | | | | Undetermined |
| 4 | 1:1, 50% VE | 0.228 | | | | | 1.000 | | | | | | | | | Undetermined |
| 5 | 1:1, 10% Chol | 0.225 | 0.469 | | | | 1.000 | 2.084 | | | | 1.000 | | | | Lamellar |
| 6 | 1:1, 25% Chol | 0.209 | 0.235 | 0.411 | 0.619 | | 1.000 | 1.124 | 1.967 | 2.962 | | 1.000 | 1.749 | 2.634 | | Mixed phase (Inv. Hexagonal + X) |
| 8 | 1:1, 10% MeOL | 0.241 | 0.245 | | | | 1.000 | 1.017 | | | | 1.000 | | | | Mixed phase |
| 9 | 1:1, 25% MeOL | 0.234 | | | | | 1.000 | | | | | | | | | Undetermined |
| 10 | 1:1, 50% MeOL | 0.197 | | | | | 1.000 | | | | | | | | | Undetermined |
| 11 | 1:1, 10% PCL | 0.234 | 0.240 | | | | 1.000 | 1.026 | | | | 1.000 | | | | Mixed phase |
| 12 | 1:1, 25% PCL | 0.234 | 0.240 | | | | 1.000 | 1.026 | | | | 1.000 | | | | Mixed phase |
| 14 | 1:2, 0% | 0.235 | | | | | 1.000 | | | | | | | | | Undetermined |
| 15 | 1:2, 10% VE | 0.231 | 0.463 | | | | 1.000 | 2.004 | | | | 1.000 | | | | Lamellar |
| 16 | 1:2, 25% VE | 0.220 | | | | | 1.000 | | | | | | | | | Undetermined |
| 17 | 1:2, 50% VE | 0.204 | | | | | 1.000 | | | | | | | | | Undetermined |
| 18 | 1:2, 10% Chol | 0.222 | 0.235 | 0.407 | 0.469 | | 1.000 | 1.059 | 1.833 | 2.113 | 0.000 | 1.000 | 1.732 | 1.996 | | Mixed phase (Inv. Hexagonal + X) |
| 19 | 1:2, 25% Chol | 0.211 | 0.233 | 0.418 | 0.469 | 0.627 | 1.000 | 1.104 | 1.981 | 2.223 | 2.972 | 1.000 | 1.794 | 2.013 | 2.691 | Mixed phase (Inv. Hexagonal + X) |
| 21 | 1:2, 10% MeOL | 0.233 | 0.401 | 0.465 | | | 1.000 | 1.721 | 1.996 | | | 1.000 | 1.160 | | | Mixed phase (Lamellar + X) |
| 22 | 1:2, 25% MeOL | 0.215 | | | | | 1.000 | | | | | | | | | Undetermined |
| 23 | 1:2, 50% MeOL | 0.174 | | | | | 1.000 | | | | | | | | | Undetermined |
| 24 | 1:2, 10% PCL | 0.235 | 0.477 | | | | 1.000 | 2.030 | | | | 1.000 | | | | Lamellar |
| 25 | 1:2, 25% PCL | 0.235 | 0.470 | | | | 1.000 | 2.000 | | | | 1.000 | | | | Lamellar |
| 27 | 1:4, 0% | 0.247 | 0.430 | 0.493 | | | 1.000 | 1.741 | 1.996 | | | 1.000 | 1.147 | | | Inverse hexagonal |
| 28 | 1:4, 10% VE | 0.242 | | | | | 1.000 | | | | | | | | | Undetermiend |
| 29 | 1:4, 25% VE | 0.226 | | | | | 1.000 | | | | | | | | | Undetermined |
| 30 | 1:4, 50% VE | 0.214 | | | | | 1.000 | | | | | | | | | Undetermined |
| 31 | 1:4, 10% Chol | 0.250 | 0.496 | | | | 1.000 | 1.984 | | | | 1.000 | | | | Lamellar |
| 35 | 1:4, 25% MeOL | 0.226 | | | | | 1.000 | | | | | | | | | Undetermined |
| 36 | 1:4, 50% MeOL | 0.195 | | | | | 1.000 | | | | | | | | | Undetermined |
| 37 | 1:4, 10% PCL | 0.251 | | | | | 1.000 | | | | | | | | | Undetermined |

### S.3 Additional SAXS analysis

Because the trends observed in MQ water regarding individual LC-phase characteristics were conserved across media types, we present the results of those experiments here in the SI. The first section contains polys analogous to **Figure 2A-D** to visualize co-core specific trends with respect to weight fraction and charge ratio. The second section presents and discusses figures analogous to **Figure 2E&F** to further prove the observed minimal effects of charge ratio and aqueous environment on d-spacing.

#### *S3.1 d-spacing analysis for liquid crystalline nanocarriers exposed to different environments*

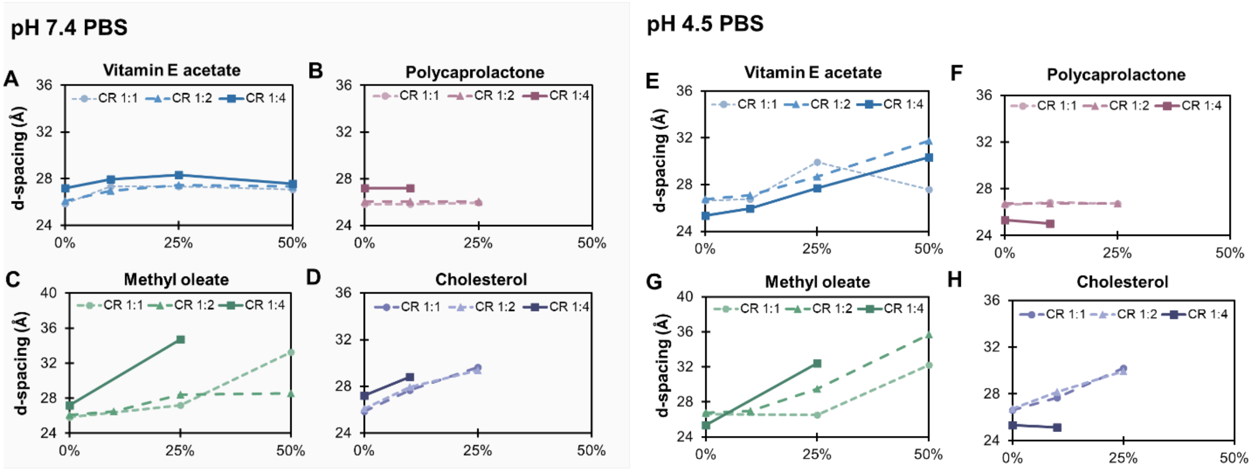

**Figure S.5:** Relationship between d-spacing and co-core weight fraction for all charge ratios and formulations including vitamin E acetate (blue), polycaprolactone (red), methyl oleate (green), and cholesterol (purple) in PBS pH 7.4 (A-D) vs PBS pH 4.5 (E-H).

#### *S3.2 Combined analysis for pH and charge ratio effects*

In the combined analyses presented below there is a shared region of the notched bar for all sample groups in both panels. Therefore, neither charge ratio or aqueous environment significantly impact the repeat distance of the LC phases.

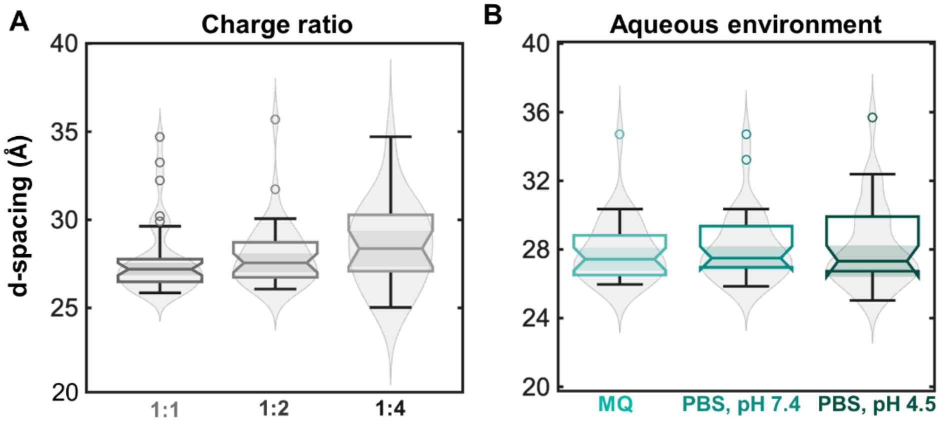

**Figure S.6:** Relationship between d-spacing and (A) charge ratio and (B) aqueous environment using notched bar charts overlayed onto violin plots.

### S.4 Development of the *in vitro* release assay protocol

#### *S4.1 Effects of CHAPS on LCNCs*

In order for CHAPS to function effectively as a hydrophobic sink, LCNCs should remain size stable in the release medium over the first few hours. As drug release becomes significant, it can be expected that morphological changes will occur. LCNCs were exposed to 10 mM CHAPS at pH 4.5 for 3 hours. After exposure, size, PDI, and surface charge were measured using the methodology reported in the manuscript and are reported in **Figure S.7**.

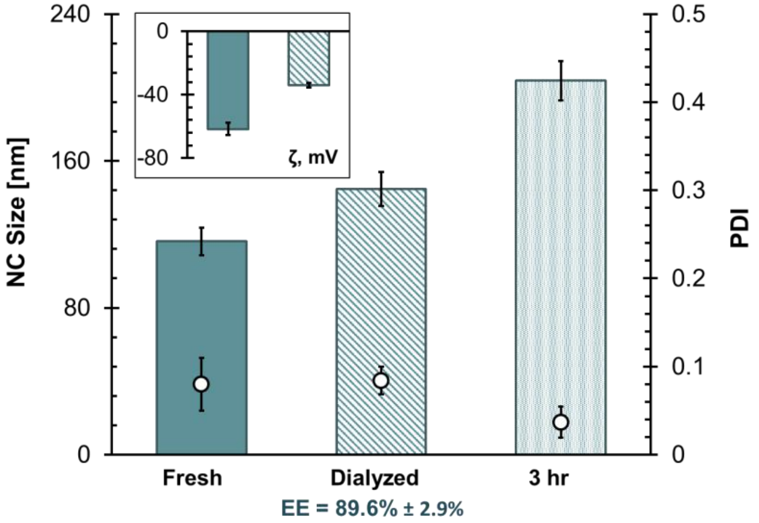

**Figure S.7**: 1:4 PMB:OL LCNCs without a co-core fresh (n = 4), dialyzed (n = 4), and after 3 hours in the release medium (n = 2). Across conditions LCNC size increased slightly, but not beyond what was observed for NCs in PBS alone (**Figure S.7**). LCNCs without a co-core are the least robust to changes in aqueous environment and were therefore selected for this testing. Zeta potential in the release media was not measured due to excess ionic strength imparted by PBS.

#### *S4.2 PMB extraction method development*

To quantify the amount of PMB in the dialysis reservoir overtime an in-house extraction protocol was developed. Validation of this protocol involved comparing extracted samples of: free PMB, free PMB processed through FNP, fresh LCNCs and dialyzed LCNCs to one another and un-extracted samples of the free drug at the fresh and FNP conditions. The results of this are provided below in **Table S.6**.

**Table S.6**: Results of PMB extraction protocol development.

| HPLC Sample Description | [Source] Theoretical | [Vial] Theoretical | [Vial] Actual | % Vial (Efficiency) | [Source] Actual | % Source (Normalized) |
|---|---|---|---|---|---|---|
| Feed stock | 5 | 0.5 | 0.49 | 98.51% | 4.926 | 99% |
| PMB -> FNP | 0.5 | 0.25 | 0.26 | 104.61% | 0.523 | 105% |
| Extracted Feed | 5 | 0.535 | 0.5 | 105.57% | 5.648 | 113% |
| Extracted PMB -> FNP | 0.5 | 0.214 | 0.20 | 93.73% | 0.497 | 99% |
| Extracted NCs | 0.5 | 0.214 | 0.19 | 88.37% | 0.493 | 93.79% |
| Dialyzed NCs (n=3) | 0.5* | 0.214 | 0.15 ± 0.01 | 70% ± 3% | 0.39 ± 0.01 | 79% ± 4% |

From this screen, it could be determined that the extraction efficiency of this process is 93.79%. Therefore, a multiplication factor of 1.06 was applied to all extraction results to compensate for any processing inefficiencies. Removal of this factor does not alter any trends in the dataset.

##### *S4.3 Preliminary work using dialysis tubing with a 6-8 kDa MWCO tubing*

To validate the working PMB extraction protocol without consuming the Float-a-Lyzer tubes reserved for the study, a series of tests (n = 3) were performed using traditional dialysis tubing. This smaller screen compared free PMB to 1:1 and 1:4 PMB:OL LCNCs without a hydrophobic co-core in PBS at pH 4.5 and is presented in the main text (Figure 6C). A reduction in MWCO impeded the transport of free PMB across the membrane significantly compared to the 100 MWCO pore size utilized in the main text (Complete transport in 24 hr vs 6 hr respectively). In both cases, the release from the LCNC was different and slower than what was observed for the free drug, indicating that the LCNCs control the release of the API and that the difference between formulations is more apparent when using a smaller MWCO tubing.

##### *S4.4 Preliminary data investigating co-core effects*

Prior to conducting the large screen, effects of co-loading PMB:OL complexes with a hydrophobic material were probed using the same methodology described above. This initial test examined trends in LCNC size, PDI, and release rate. PMB:OL LCNCs at a 1:4 charge ratio with and without a 50% VEAc co-core were manufactured for this experiment. This proof-of-concept pilot study revealed differences in LCNC release and size evolution over of 48 hours. These results supply the basis for the main investigation reported in the main text. The condition of pH 4.5 was isolated in these preliminary experiments as release at pH 7.4 was hypothesized (and proven in the main text) to be minimal over this window. The results of this pilot study are provided in **Figure S.8**.

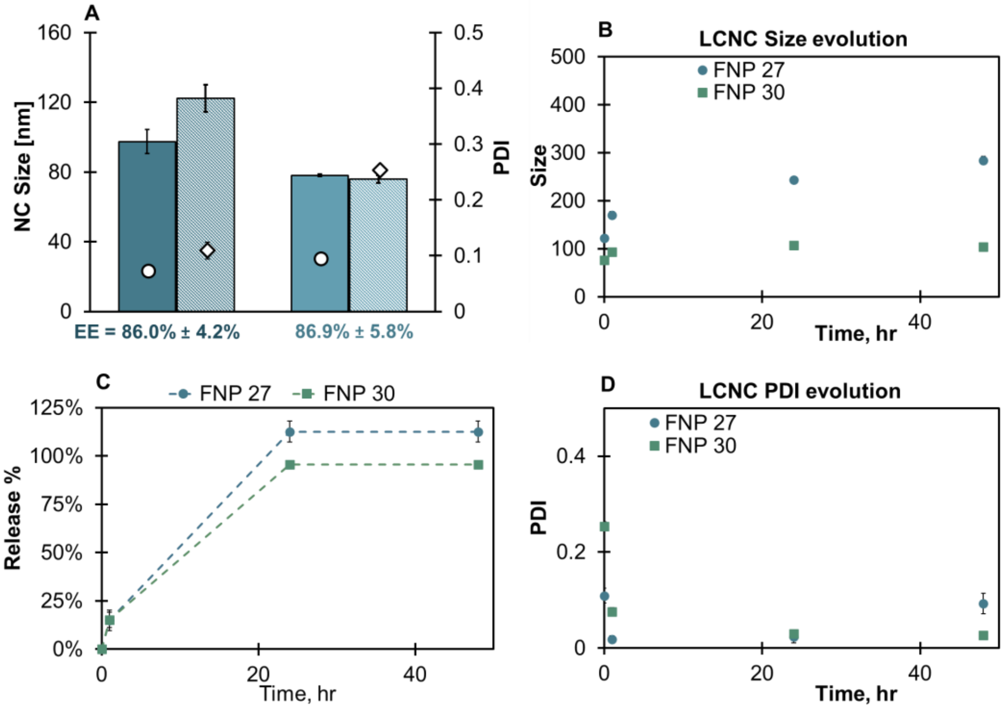

**Figure S.8:** (A) LCNC properties pre and post dialysis at a fixed 1:4 charge ratio with (FNP 30) and without (FNP 27) a hydrophobic co-core of vitamin E acetate at 25% by weight.

*S4.5 CHAPS validation*

CHAPS is a zwitterionic low-aggregation number surfactant reported to form micelles at low concentrations (8 mM). These properties make CHAPS an ideal candidate as a hydrophobic sink as they are able to transit a 100 kDa membrane. The membrane is critical in this work to separate encapsulated drug (within the LCNCs) from the bulk release media facilitating analysis of drug release over time. Solution properties such as pH and ionic are known to influence the CMC of surfactant. Therefore, to validate that the protocol functions as hypothesized a pilot study was conducted at a concentration of 10 mM CHAPS. If CHAPS can form micelles under these conditions as theorized, the release kinetics for a given formulation should differ under each condition. The LCNCs reported in **Figure S.7** were utilized in this test. Two replicates at each condition were performed for both the LCNCs and a free drug control.

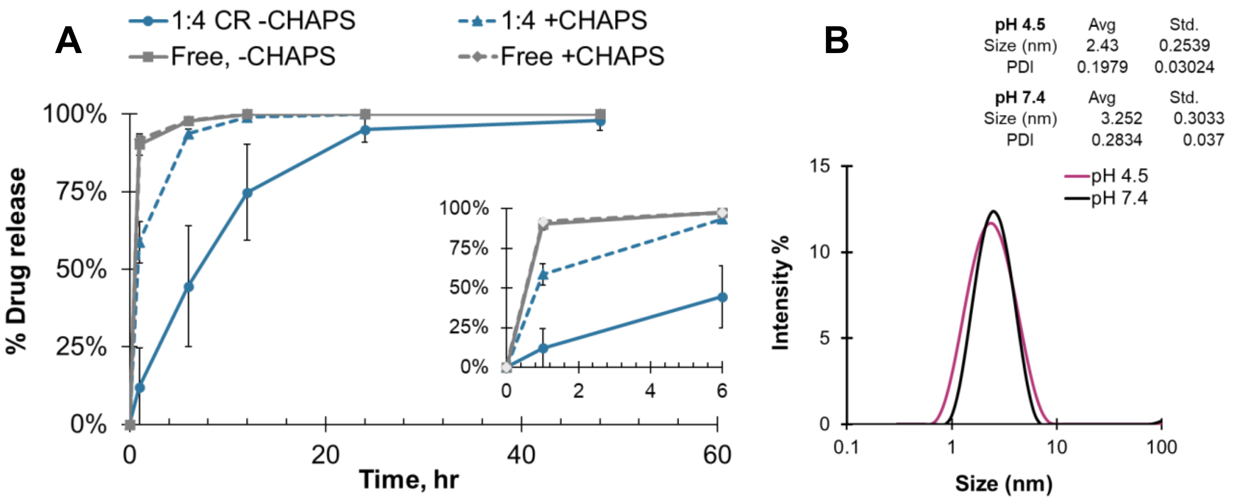

**Figure S.9**: (A) Results of free drug controls and LCNCs at a 1:4 CR without a co-core in PBS at pH 4.5 with vs without CHAPS. The inset is the same data set over the first 6 hours to visualize initial differences in release rate. (B) Dynamic light scattering analysis of CHAPS micelles.

This pilot study revealed several important factors that were applied to the overall experimental design of the main study. The first of which is that free PMB diffusion across the membrane is not influenced by the hydrophobic sink. This finding is in line with expectations of the release or diffusion of a hydrophilic drug into the hydrophilic bulk. The second key finding is that, in line with the hypothesis and many literature reports investigating drug release, the presence of CHAPS aids in the release of PMB from LCNCs. The free drug control reported here is not the same control from the main text, that control was performed using the Float-a-Lyzer devices while this test was performed using 100 kDa MWCO dialysis tubing and clamps. Notably, there is no difference in the release rates, consistent with diffusion dependance on membrane pore size.

*S4.6 Formulation parameters for media effect screen*

**Table S.7:** Summary of formulation parameters for each of the key formulations taken through the release media screen. Characterization data provided in **Figure S.12.**

| Formulation | Organic feed | Aqueous feed |
|---|---|---|
| 1:1 PMB:OL, 0% CC | 5 mg/mL PCL-b-PEG<br>5.85 mg/mL NaOL | 5 mg/mL PMB |
| 1:1 PMB:OL, 25% VE | 5 mg/mL PCL-b-PEG<br>5.85 mg/mL NaOL<br>3.16 mg/mL VEAc | 5 mg/mL PMB |
| 1:1 PMB:OL, 25% PCL | 5 mg/mL PCL-b-PEG<br>5.85 mg/mL NaOL<br>3.16 mg/mL PCL | 5 mg/mL PMB |
| 1:4 PMB:OL, 0% CC | 5 mg/mL PCL-b-PEG<br>23.4 mg/mL NaOL | 5 mg/mL PMB |
| 1:4 PMB:OL, 25% VE | 5 mg/mL PCL-b-PEG<br>23.4 mg/mL NaOL<br>9.49 mg/mL VEAc | 5 mg/mL PMB |
| 1:4 PMB:OL, 25% PCL | 5 mg/mL PCL-b-PEG<br>23.4 mg/mL NaOL<br>9.49 mg/mL PCL | 5 mg/mL PMB |

#### S.5 HPLC calibration curve data

The calibration curves utilized to analyze the APIs in this study were formed under the conditions that samples were to be analyzed.

##### *S5.1 Polymyxin B calibration curve*

The PMB calibration curve was generated using samples of free drug subject to the extraction protocol. Since PMB is a heterogenous mixture of polymyxins B1 and B2, there are multiple peaks to integrate per sample. To normalize this across runs, peaks from 4.4 min to 5.4 min were considered. This window is consistent with what is observed for the obtained lot of PMB in water before and after filtration. The calibration curve utilized to analyze the release data is provided as **Figure S.10**.

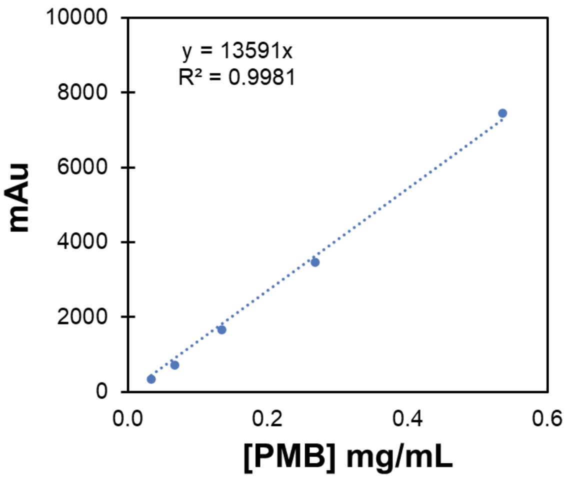

**Figure S.10:** Calibration curve, equation, and $R^2$ for extracted polymyxin B.

##### S.5.2 Vitamin E acetate calibration curve

The VEAc calibration curve was generated using VEAc dissolved in THF. The resolution of the VEAc peak in THF is poor compared to VEAc in THF. However, THF was the only solvent that was able to extract VEAc with an efficiency >70%. To normalize this across runs, peaks from 3.5 min to 6.4 min were considered. VEAc extraction trials, controls and calibration curve data are provided in **Figure S.11**.

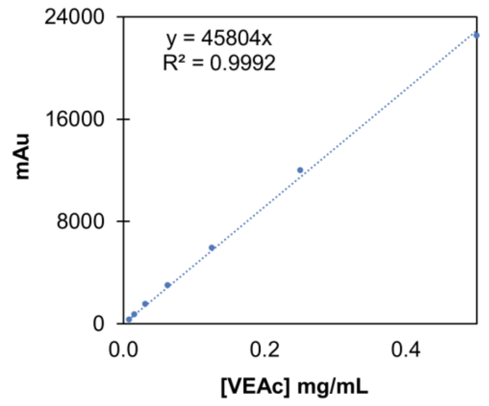

| Sample | [Stock] | [Ext] | mAu | [Sample] | %Eff | |
|---|---|---|---|---|---|---|
| 1-UF | 0.95 | 0.407143 | 19463.9 | 0.40183 | 98.69% | |
| 1-F | 0.95 | 0.407143 | 19507.8 | 0.402747 | 98.92% | |
| 2-UF | 0.35 | 0.15 | 7166.79 | 0.144815 | 96.54% | |
| 2-F | 0.35 | 0.15 | 7197.43 | 0.145456 | 96.97% | |
| 28 | 0.35 | 0.15 | 1752.6 | 0.031657 | 21.10% | |
| 29 | 0.95 | 0.407143 | 1431.91 | 0.024954 | 6.13% | |
| 28-2 | 0.35 | 0.35 | 11961.3 | 0.245022 | 70.01% | ACN (100%) |
| 29-2 | 0.95 | 0.475 | 14855.6 | 0.305514 | 64.32% | ACN (100%) |
| 1-F. THF | 0.95 | 0.95 | 44138.7 | 0.917543 | 96.58% | |
| 2-F. THF | 0.32 | 0.32 | 15295.7 | 0.314713 | 98.35% | |
| 28-1-1 | 0.32 | 0.32 | 13229.8 | 0.271535 | 84.85% | |
| 28-1-1.5 | 0.32 | 0.213333 | 9172.21 | 0.186729 | 87.53% | |
| 28-0.5-2 | 0.32 | 0.16 | 6955.22 | 0.140393 | 87.75% | |
| 29-1-1 | No sample injected | | | | | |
| 29-1-1.5 | 0.95 | 0.633333 | 27689 | 0.573737 | 90.59% | |
| 29-0.5-2 | 0.95 | 0.475 | 20029.2 | 0.413645 | 87.08% | |

**Figure S.11:** Calibration curve and extraction data for VEAc.

Trials that resulted in extraction efficacy < 20% applied the same methodology utilized to extract PMB from LCNCs. The excess water prohibited proper solubilization of VEAc from LCNCs despite LCNC structure disruption. Therefore, aliquots were dried and resuspended in 100% solvent (first ACN, then THF). This workflow yielded more optimal extraction results and was employed in the key experiments presented in the main text.

### S.6 Nanocarrier characteristics and performance

#### *S6.1 Characteristics of liquid crystalline nanocarrier formulations*

Size, PDI, and encapsulation efficiency for the nanocarrier formulations in the main text are presented here. Zeta potential, as discussed in the main text and in **Section S6.2**, was found to be largely independent of co-core composition and is instead reported for LCNCs made at each charge ratio. The ripening of the nanocarriers in **Figure S.12A** for the 1:1 PMB:OL formulations is expected. These formulations require inclusion of a co-core for consistency in stability over dialysis. As show in the main text, the same formulation reproduced for the mucus diffusion work ripened only to 250 nm. Throughout the study and preliminary work, the variability in the extent to which the 1:1 PMB:OL formulation ripens is 1) consistent (+200-400 nm) and 2) is attributable to formulation instabilities which can be mitigated through including a hydrophobic co-core or increasing the charge ratio. Control over the dialysis process via pH or ionic strength could also mitigate this ripening, however, to maintain parity across all formulations the dialysis process was kept consistent.

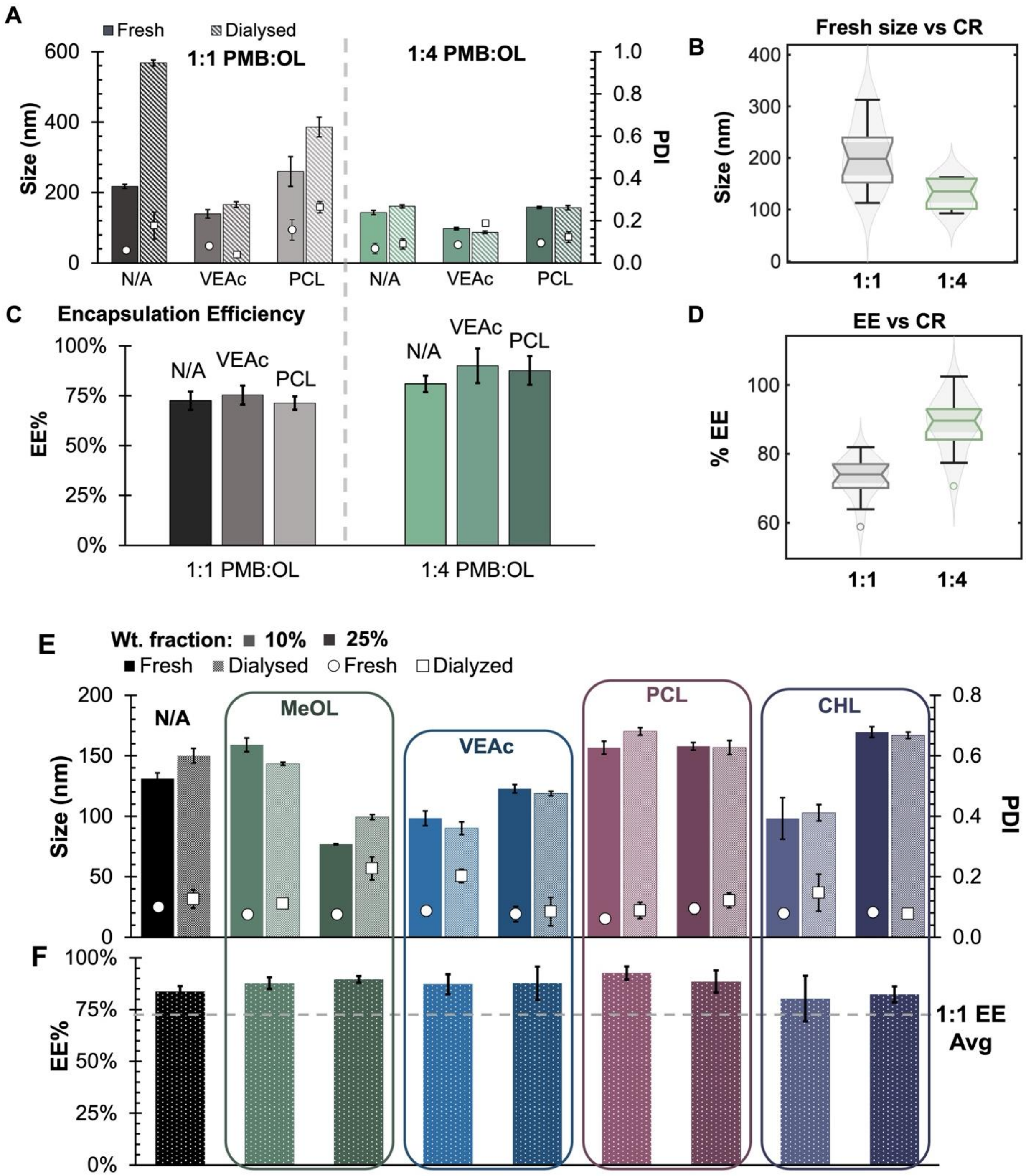

**Figure S.12:** (A) Size and PDI and (C) encapsulation efficiency of six key formulations. (B&D) Notched box plots overlayed onto violin plots to highlight relationship between LCNC formulation parameters and physicochemical properties considering the data distribution for fresh suspensions. In this figure, all co-core weight fractions are fixed to 25%. Error bars represent the standard deviation of n = 6 replicates. (D) Size, PDI, and (E) encapsulation efficiency of LCNC formulations co-encapsulating PMB with each of four different hydrophobic co-cores with the no co-core sample plotted as a control. All formulations were formulated at a fixed 1:4 charge ratio and error bars represent the standard deviation of n = 3 runs. Bar shade correlates with weight fraction where the lighter bar (left side) is 10% and the darker bar (right side) is 25% co-core by mass.

##### S6.2 Zeta potential analysis for liquid crystalline nanocarriers

Nanocarrier surface charge is sensitive to the buffer in which the measurement is made, particularly for pH sensitive systems. This is particularly true for nanocarriers containing hydrophobic counterion or any other pH sensitive materials. Measurement of sample zeta potential in **Section S.4** was done using 20 mM NaCl. Because this solution is unbuffered, the pH over time is uncontrolled potentially compromising the experimental determinations of zeta potential generating misleading data. After screening buffer candidates using nanocarrier components as controls, pH 7.2 HEPES supplemented with KCl was determined to be the optimal choice (**Figure S.13**).

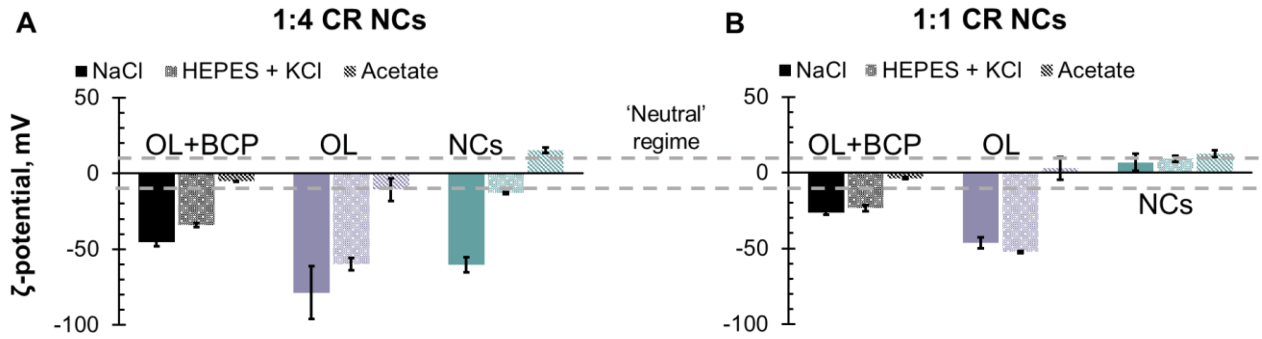

**Figure S.13:** Zeta potential readings for LCNCs, counterion only, and PCL-PEG + counterion samples formulated via FNP at a 1:4 (A) vs 1:1 (B) charge ratio.

##### S6.3 Similarity factor analysis, $f_2$

The similarity factor or $f_2$ is an FDA-recognized statistical test classically employed to determine if two dissolution kinetics curves are similar to one another [1, 2]. The guidance for employing this test states that the data set should contain low variability, at least three shared time points, and include at least no more than one data point > 85% release. The statistical power increases with the number of replicates (12 is the FDA recommended number). In this work, all criteria are satisfied except for the number of replicates. However, given that the replicates acquired have low variability we apply **Equation S.1**, below, to calculate the $f_2$ value as a way to confirm observations about the effects of different media on a given formulation in Figure 4 in the main text and **Figure S.14** here.

$$\boldsymbol{Eqn\,S.1}:\ f_2 = 50\ \times log_{10}\left\{\left[1+\frac{1}{n}\sum_{t=1}^{n}(R_t - T_t)^2\right]^{-0.5} \times 100\right\}$$

In this equation, n is the number of timepoints, $R_t$ = the average value for the percent released in the reference curve scaled from 0-100 at time t, and $T_t$ = the average value for the percent released in the test curve scaled from 0-100 at time t. If the resulting $f_2$ is ≥50, the curves are considered to be not significantly different from one another. Conversely, if the $f_2$ is <50, the curves are different from one another with the magnitude scaling as the value decreases [2].

*S6.4 Release kinetics co-core data, alternate plots by media type and charge ratio*

**Figure S.14** is complementary to **Figure 3** in the main text. The trends down each column recapitulate what was observed in **Figure 3** further supporting that the effects of changing the release media are conserved across all formulation types. This is to say that switching the media type would not result in a change in the release kinetics hierarchy.

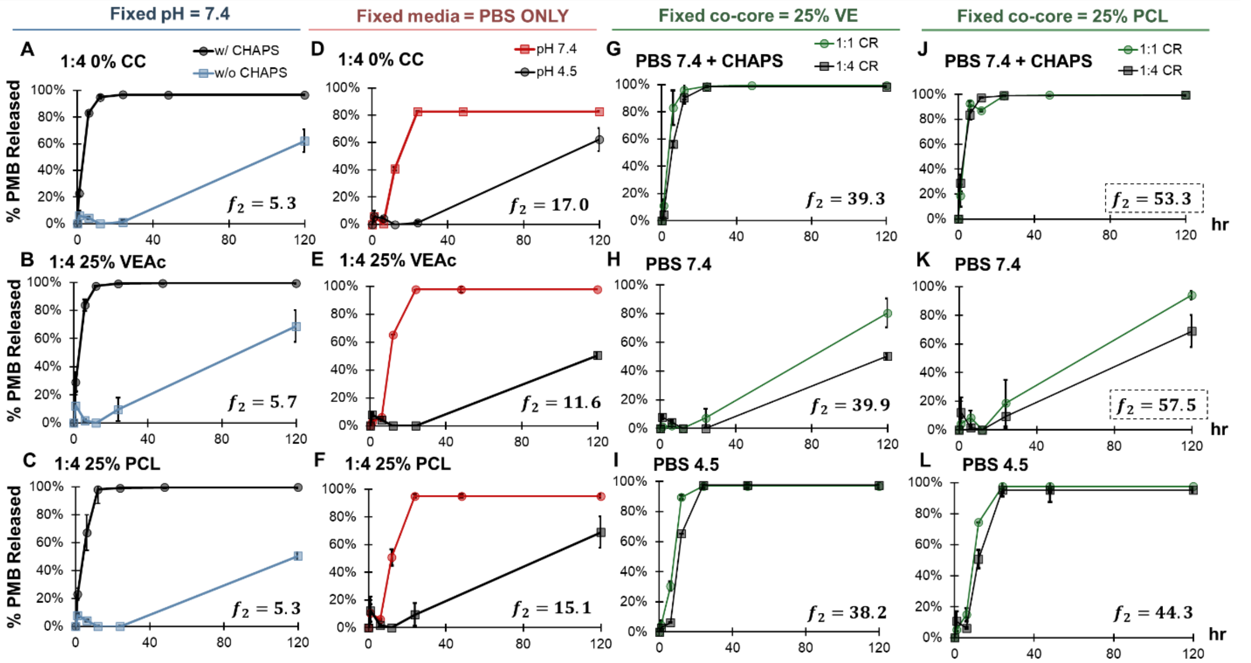

**Figure S.14:** Pairwise comparisons at a fixed pH comparing media containing CHAPS vs without (A-C), in the absence of CHAPS comparing pH (D-E), and at all three media conditions but at fixed co-core type and weight fraction (G-L). Error bars represent the standard deviations for n = 3 separate tests. $f_2$ values were determined using the methodology described in **S.6.3.** A dotted box was included in panels J&K to indicate the only set of curves that are similar to one another according to the analysis.

##### *S6.5 Release kinetics behavior of co-encapsulated hydrophobic payload VEAc*

The release kinetics of VEAc from LCNCs exhibits a significant dependance on the presence of the hydrophobic sink and slight dependance on drug loading. **Figure S.15A&B**, compares the release kinetics of VEAc at a 25% weight fraction from LCNCs formulated at a 1:1 vs a 1:4 CR in the presence and absence of CHAPS. **Figures S.15C&D** isolate LCNCs at a 1:4 CR to probe the effects of weight fraction on VEAc release. All pairwise comparisons were made using the methodology detailed in **Section S6.4**. However, only the difference in VEAc release at t = 6 hr comparing 10% to 25% wt fraction was significant. Statistical evaluation was not performed comparing the release of VEAc in the presence/absence of CHAPS as the amount of VEAc remained the same over time, indicating that no release occurred over the experimental period. This result as expected because without a hydrophobic sink, there is no driving force for VEAc to partition out of the LCNC core. Data is plotted to 24 hrs to showcase the time points of interest.

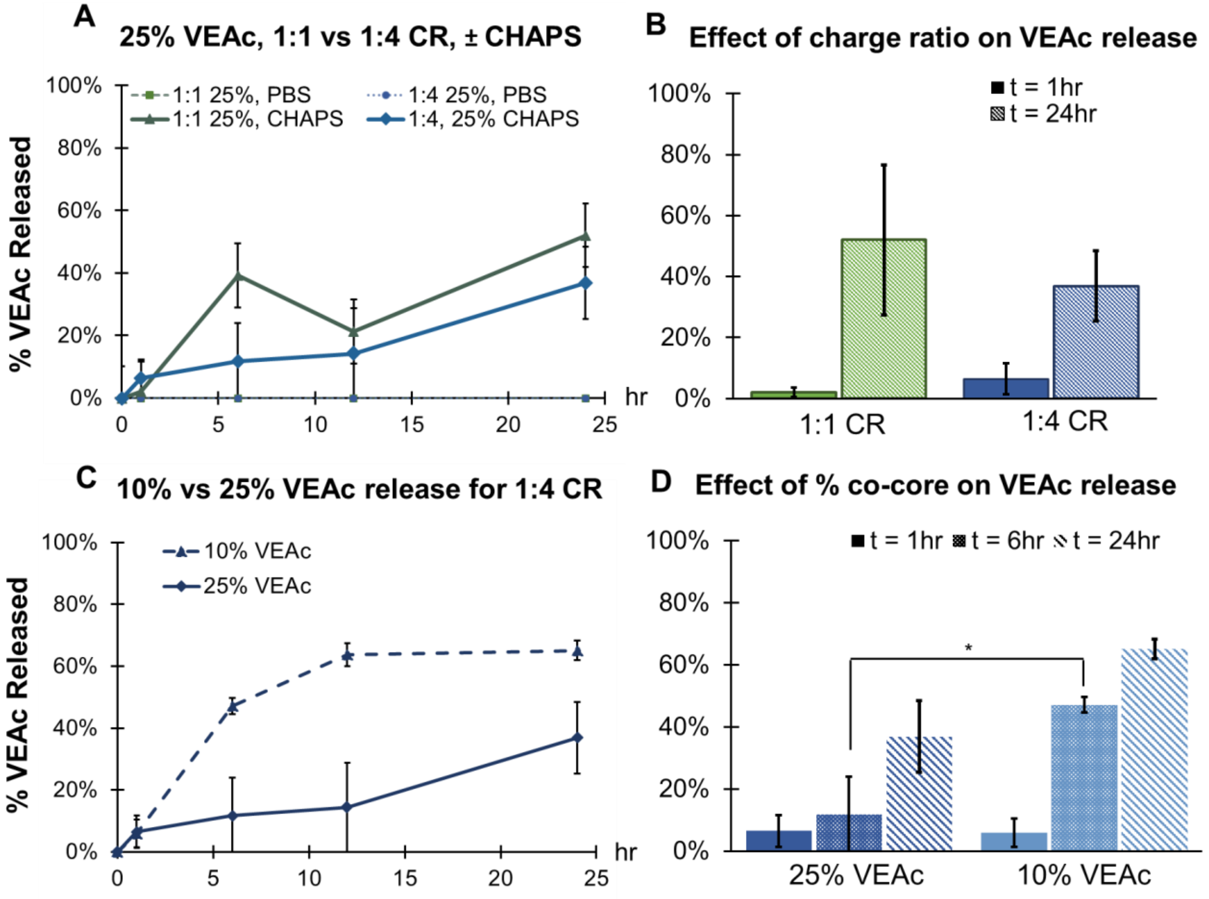

**Figure S.15:** (A) Release kinetics profiles of LCNCs at a 1:1 vs 1:4 charge ratio with VEAc at a 25% weight fraction in the presence and absence of CHAPS. (B) Bar chart isolating the release at a 1:1 vs 1:4 charge ratio in the presence of CHAPS at t = 1 hr vs t = 24 hr. (C) LCNCs at a 1:4 CR containing either a 10% (dotted line) or a 25% VEAc co-core (solid line) by mass. (D) Bar chart isolating key timepoints from the release kinetics curve in (C) Error bars represent the standard deviations for n = 3 separate tests.

*S6.6 Tabulated summary of co-core physicochemical and thermodynamic parameters*

To establish if there is one predictive parameter that may be leveraged to predict release behavior, a variety of properties were examined and summarized in the table below. LogP values were determined according to the provided references, the solid states at room temperature were determined through observation of purchased material, and the Hansen distances were calculated using the methodology below.

**Table S.8:** Summary of co-core and hydrophobic counterion specific parameters. Where formulation specific, data refers to the 25% wt. fraction.

| Material | LogP | State | $R_a$ $(\frac{J}{cm^3})^{1/2}$ | d-spacing (Å) | %Release$_{6hr}$ |
|---|---|---|---|---|---|
| Oleate (ion paired) | 4.9 [3] | Solid | - | 25.3 | 83.1 ± 1.3% |
| Methyl oleate | ~ 7.5 [4] | Oil | 13.4 | 34.7 | 49.1 ± 2.8% |
| Polycaprolactone (2K) | 7.2 [5] | Solid | 36.1 | 25.7 | 83.7 ± 3.4% |
| Cholesterol | ~ 8.8 [6] | Solid | 33.1 | 29.4 | 65.8 ± 6.3% |
| Vitamin E acetate | 10.4 [5] | Viscous oil | 3.4 | 28.3 | 72.2 ± 2.0% |

Evaluating the thermodynamic favorability of the interaction between the hydrophobic counterion complex and each hydrophobic co-core material was done by calculating the Hansen distance (**Equation S.2**) [7].

$$\boldsymbol{Eqn\ S.2}: \mathrm{R}_a = \sqrt{4(\delta\mathrm{D}_1 - \delta\mathrm{D}_2)^2 + (\delta\mathrm{P}_1 - \delta\mathrm{P}_2)^2 + (\delta\mathrm{H}_1 - \delta\mathrm{H}_2)^2}$$

In this equation, $R_a$ is the Hansen distance, 1 refers to the value for the reference phase (oleate) and 2 refers to the sample phase (one of four hydrophobic co-cores). Each $\delta$ represents one of the Hansen solubility parameters where $\delta$D refers to the dispersion component (captures the effects of induced dipole-induced dipole interactions), $\delta\mathrm{P}$ refers to the polar component (captures the contribution of polar groups), and $\delta\mathrm{H}$ is the hydrogen bonding component (captures the effects of hydrogen bonding). All parameters have units of $(\frac{J}{cm^3})^{1/2}$.

The Hansen parameters ($\delta$) in **Equation S.2** were determined using the Hoftyzer-Van Krevelen equations provided below (**Equations S.3-5**) [8].

$$\boldsymbol{Eqn\ S.3}: \delta_D = \frac{\sum F_{di}}{V}$$

$$\boldsymbol{Eqn\ S.4}: \delta_P = \frac{\sum {F_{pi}}^2}{V}$$

$$\boldsymbol{Eqn\ S.5}: \delta_H = \frac{\sum E_{hi}}{V}$$

The sum of $F_{di}$, ${F_{pi}}^2$, and $E_{hi}$ were respectively determined using Table 4 “Solubility Parameter I: Component Group Contributions” in the Polymer Handbook, 4th ed. By Brandrup, Immergut & Grulkable on page 686 along with the values commonly adapted in polymer science literature [9]. The contribution of PMB is constant across materials and was not accounted for and the selected model does not account for well salts, thus a -COO- group contribution was utilized as the terminal group for sodium oleate. In **Eqns S.3-5**, V is the molar volume was determined via **Equation S.6.**

$$\textbf{Eqn S.6}: V = \frac{M}{\rho}$$

In **Equation S.6**, $\rho$ = the density of each material at room temperature (g/cm$^3$) and $M$ = the molecular weights of each material (g/mol) as provided by the manufacturer unless a reference is listed (**Table S.9**). It is important to note that including the contribution of polymyxin B would increase the applicability of such thermodynamic favorability interactions. Furthermore, as charge ratio increases, the model would need to account for the various charge states of sodium oleate (neutral when complexed to PMB, excess would be charged). Such analyses would require complex computational tools and should be the focus of a dedicated future work.

**Table S.9**: Data used to determine the molar volume using **Eqn S.6**

| Material | MW (g/mol) | Density (g/cm$^3$) |
|---|---|---|
| Oleate (ion paired) | 296.49 | 0.874 |
| Methyl oleate | 472.74 | 0.955 [4] |
| Polycaprolactone (2K) | 2000 | 1.071 |
| Cholesterol | 386.65 | 1.067 |
| Vitamin E acetate | 282.46 | 0.8925 [10] |

#### *S6.7 Statistical analysis of release kinetics data – Figure 6*

To more accurately assess the influence of co-core physicochemistry and weight fraction on drug release, a three-way fixed effects ANOVA was run with a *post hoc* Tukey HSD. The general model for this analysis is provided below as equation S.7.

$$\textbf{Eqn S.7}: Y_{ijkl} = \mu + \alpha_i + \beta_j + \gamma_k + (\alpha\beta)_{ij} + (\alpha\gamma)_{ik} + (\beta\gamma)_{jk} + (\alpha\beta\gamma)_{ijk} + \epsilon_{ijkl}$$

In this model, $Y_{ijkl}$ represents a specific observation (fraction polymyxin B released) and represent the i,j,k are levels of factors alpha, beta, gamma and l represents the replicate for that specific combination of levels where: $\alpha_i$ = main effect of co-core type, $\beta_j$ = main effect of co-core mass fraction, and $\gamma_k$ = the main effect of time.

The results of the analysis for the main effects of co-core type and weight fraction reveal that the co-core type is significant except at 12 h. At 1 and 6 h, the type is significant at both weight fractions. The final pairwise comparisons determined using a *post hoc* Tukey HSD are presented in the table below using a compact letter display.

**Table S.10**: Compact letter display for the main effects of co-core type averaged across all weight levels.

| Time | CHL | MeOL | PCL | VEAc |
|---|---|---|---|---|
| 1 h | a | a | a | a |
| 6 h | a | a | b | b |
| 12 h | a | a | a | a |

Performing the same analysis for co-core type, it was found that the weight fraction of co-core only has a significant effect at time 12 h. Ultimately, across the presented three time points and the four types of co-core the main effects of weight fraction are not significant ($F_{1,48}$ = 3.13, p-value = 0.0830). However, there are only two levels of mass fraction (10% and 25%), so the df for the numerator of the F-statistic is only 1, severely limiting the statistical power of the test. It is also important to note that the interactions between these factors is significant $F_{1,48}$ = 24.80, p-value = <0.0001.

The second analysis conducted involved comparing the release kinetics of each formulation to that of the free drug and formulations without a co-encapsulated payload. For this analysis, a two-way ANOVA where co-core type and weight have been grouped into the same factor was performed applying the same assumptions discussed and evaluated above. The results of this analysis show that there is an observed effect for the interaction between co-core type and time (F-value = 9.97, p-value = <0.0001). The pairwise comparisons for the *post hoc* Tukey HSD are provided below in table **S.11 (**vs Free drug) and **S.12** (vs No co-core control).

**Table S.11:** Pairwise comparisons against the free drug.

| Time | No CC | CHL10 | CHL25 | MeOL10 | MeOL25 | PCL10 | PCL25 | VE10 | VE25 |
|---|---|---|---|---|---|---|---|---|---|
| 1 h | <.0001 | <.0001 | <.0001 | <.0001 | <.0001 | <.0001 | <.0001 | <.0001 | <.0001 |
| 6 h | 0.1123 | <.0001 | <.0001 | <.0001 | <.0001 | <.0001 | 0.1805 | <.0001 | <.0001 |
| 12 h | 1.0000 | 1.0000 | 0.8778 | 1.0000 | 0.6221 | 1.0000 | 1.0000 | 1.0000 | 0.8910 |

†All pairwise comparisons: $F_{1,60}$, Tukey-adjusted p-values shown.

**Table S.12:** Pairwise comparisons against the 0% CC samples.

| **Time** | **Free** | **CHL10** | **CHL25** | **MeOL10** | **MeOL25** | **PCL10** | **PCL25** | **VE10** | **VE25** |
|---|---|---|---|---|---|---|---|---|---|
| 1 h | <.0001 | 0.0085 | 1.0000 | 1.0000 | 0.0002 | 0.9980 | 0.9868 | 1.0000 | 0.9999 |
| 6 h | 0.1123 | <.0001 | 0.0008 | 0.0004 | <.0001 | 0.2153 | 1.0000 | 0.4752 | 0.2343 |
| 12 h | 1.0000 | 1.0000 | 0.9988 | 1.0000 | 0.9700 | 1.0000 | 1.0000 | 1.0000 | 0.9991 |

†All pairwise comparisons: $F_{1,60}$, Tukey-adjusted p-values shown.

#### S.7 Efficacy studies against a bacterial pathogen

##### *S7.1 Effects of release medium on bacterial growth*

To facilitate release of PMB over timescales relevant to the *in vitro* assay the nanocarrier suspensions were diluted in release media (PBS pH 4.5) to achieve 2x desired final concentration in each well. Prior to implementing this strategy, bacterial viability in each of the media was evaluated using $OD_{600}$ measurements over time after dosing 100 uL of media onto 100 uL of bacteria diluted to a concentration of ~$5x10^6$ cfu/mL. The results of this viability study are presented in **Figure S.16** and reveal that the bacteria are sensitive to CHAPS but remain viable in PBS at either pH.

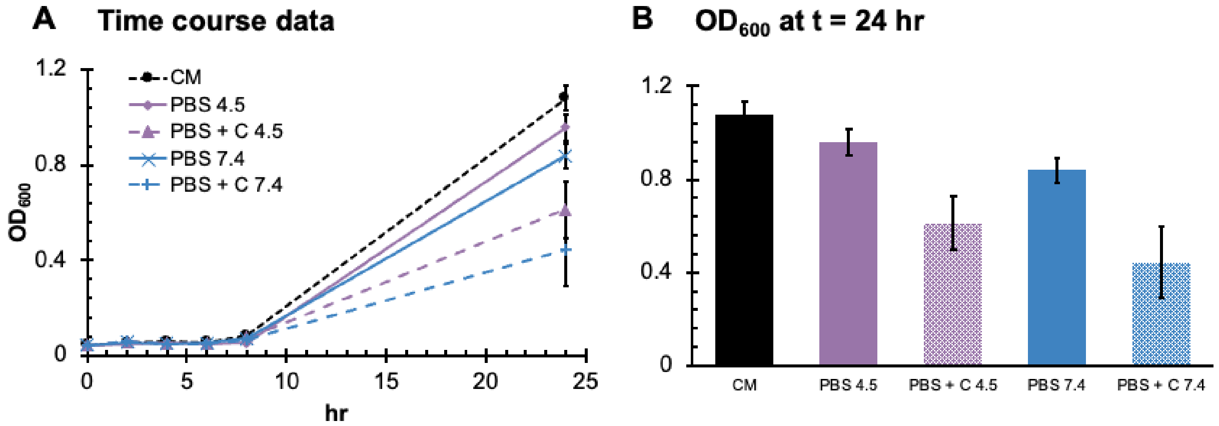

**Figure S.16**: (A) Time course study monitoring bacteria growth every two hours for up to 8 hrs then after 24 hrs. (B) Bar charts isolating the 24 hr time point. CM = control of bacteria in the culture media, C = CHAPS. Error bars represent the standard deviation of 12 wells.

##### *S7.2 Initial minimum inhibitory concentration determination*

Before experimental work with formulations was initiated, multiple free drug trials were performed to establish a baseline and develop a working protocol. The results of the free drug trials are presented below. The data show consistently over different trials that the MIC of the free drug is ~0.5 ug/mL which demonstrates strong parity with literature reports.

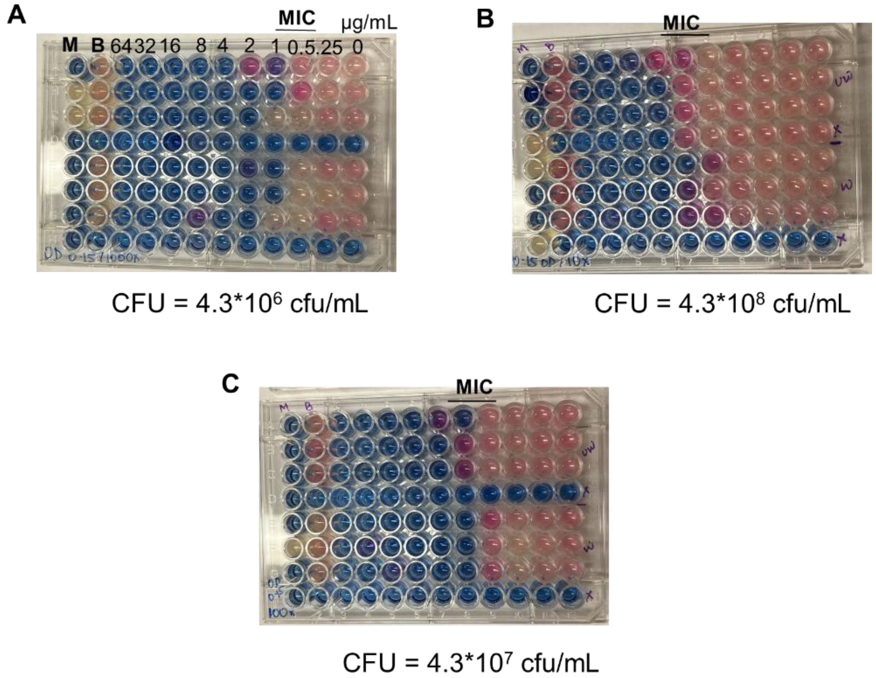

**Figure: S.17**: (A-C) Resazurin assay on free polymyxin B sulfate at 3 different bacterial concentration.

##### *S7.3 Dilution tests for streaking bacterial colonies to determine CFU*

To establish the dilution factor necessary to determine the reduction in colony forming units at the MIC and surrounding concentration a screen of treated wells vs the controls at four dilution factors was conducted. This screen investigated wells at different optical densities. **Figure S.18** contains a summary of the collected data and identified dilution factors for streaking.

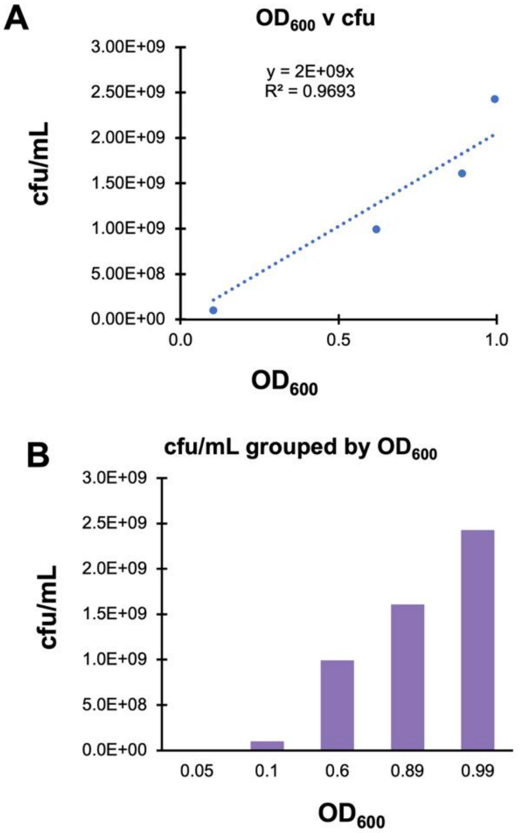

C

| Treatment | Well | OD600 | Dilution | Colonies | cfu/mL |
|---|---|---|---|---|---|
| Control | 9E | 0.891 | 1000000 | 71 | 1.42E+09 |
| Control | 9E | 0.891 | 10000000 | 7 | 1.40E+09 |
| Control | 9E | 0.891 | 100000000 | 1 | 2.00E+09 |
| Control | 9E | 0.891 | 1000000000 | 0 | - |
| Control | 8C | 0.994 | 1000000 | 164 | 3.28E+09 |
| Control | 8C | 0.994 | 10000000 | 10 | 2.00E+09 |
| Control | 8C | 0.994 | 100000000 | 1 | 2.00E+09 |
| Control | 8C | 0.994 | 1000000000 | 0 | - |
| Free drug | 6C | 0.048 | 100 | 0 | - |
| Free drug | 6C | 0.048 | 1000 | 0 | - |
| Free drug | 6C | 0.048 | 10000 | 0 | - |
| Free drug | 6C | 0.048 | 100000 | 0 | - |
| Free drug | 6B | 0.62 | 100 | lawn | |
| Free drug | 6B | 0.62 | 1000 | lawn | |
| Free drug | 6B | 0.62 | 10000 | lawn | |
| Free drug | 6B | 0.62 | 100000 | 496 | 9.92E+08 |
| NC | 6E | 0.104 | 100000 | 67 | 1.34E+08 |
| NC | 6E | 0.104 | 10000 | 324 | 6.48E+07 |
| NC | 6E | 0.104 | 1000 | lawn | |
| NC | 6E | 0.104 | 100 | lawn | |
| NC | 7E | 0.058 | 100 | 0 | - |
| NC | 7E | 0.058 | 1000 | 0 | - |
| NC | 7E | 0.058 | 10000 | 0 | - |
| NC | 7E | 0.058 | 100000 | 0 | - |

**Figure S.18:** (A&B) Relationship between cfu/mL and $OD_{600}$ based on all sampled wells averaged based on CFU counts determined from each dilution ($10^6$-$10^9$ for $OD_{600}$ > 0.7 and $10^2$ – $10^4$ for $OD_{600}$< 0.7). (C) Tabulated data summarizing the sampled well, dilution factor, and outcome of colony count determination.

*S7.4 Resazurin assay for nanocarrier formulations*

The resazurin assay output complimentary to the presented bar charts in **Figure 7** are provided below. The results of this assay reveal a consistent MIC range for all explored nanocarrier formulations at a 1:1 charge ratio and the free drug.

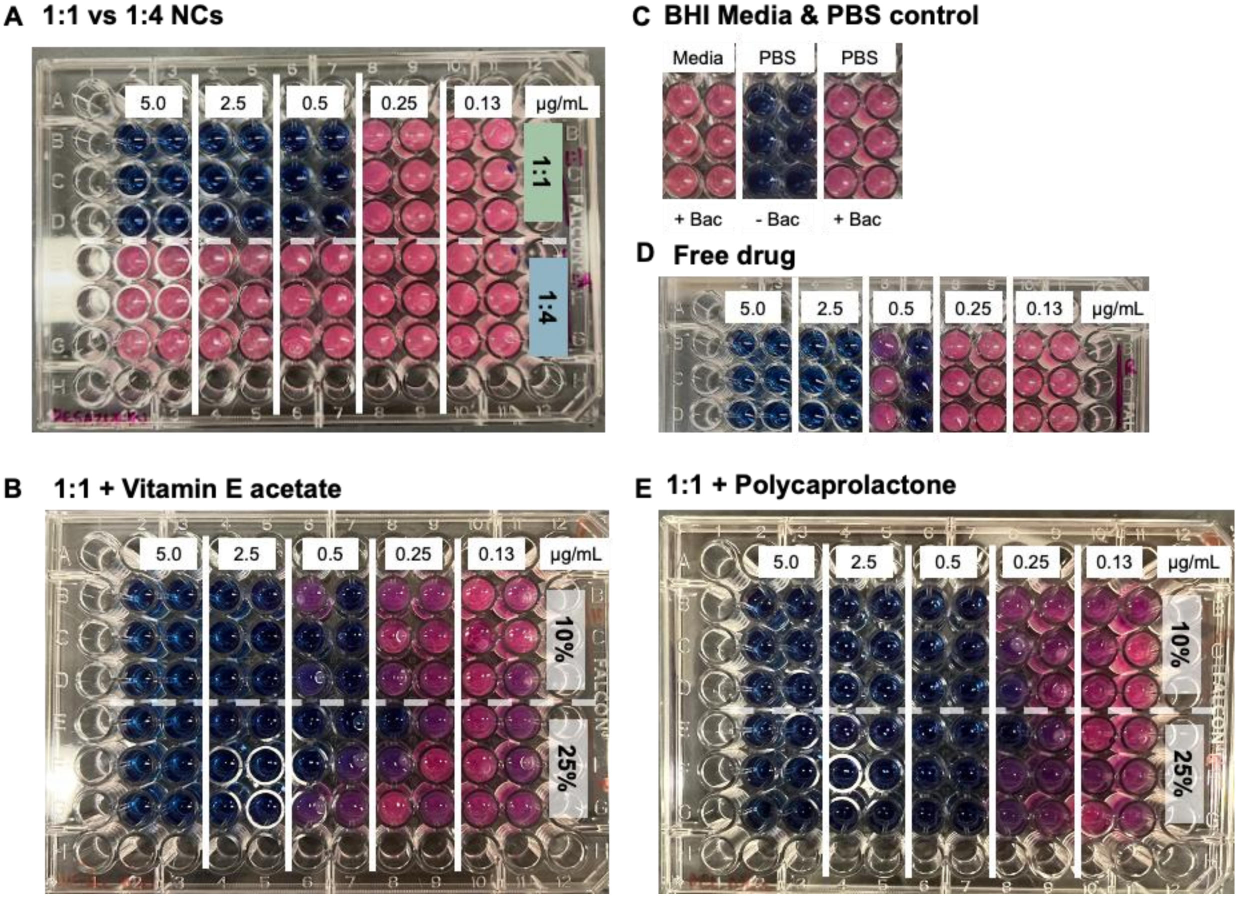

**Figure S.19:** Photographs of resazurin staining for MIC for (A) formulations at a 1:1 vs a 1:4 charge ratio, (B) formulations at a 1:1 charge ratio with 10% or 25% VEAc as the co-core, (C) controls, (D) free drug, and (E) formulations at a 1:1 charge ratio with 10% or 25% PCL as the co-core

### S.8 Isolated primary peak data presentation

#### *S8.1 Inset of scattering profiles focused on primary peak*

SAXS data for all samples in MQ water focusing on the *q* range containing the primary peak. The data below is provided to isolate trends associated with increasing charge ratio as well as increasing the weight fraction co-core for a fixed charge ratio. Data for 10% MeOL LCNCs was only collected in MQ water due to initial concerns about LCNC stability that were later determined to be unsubstantiated. As a result, there is not data for this formulation in PBS at either pH and thus the set was excluded from prior figures.

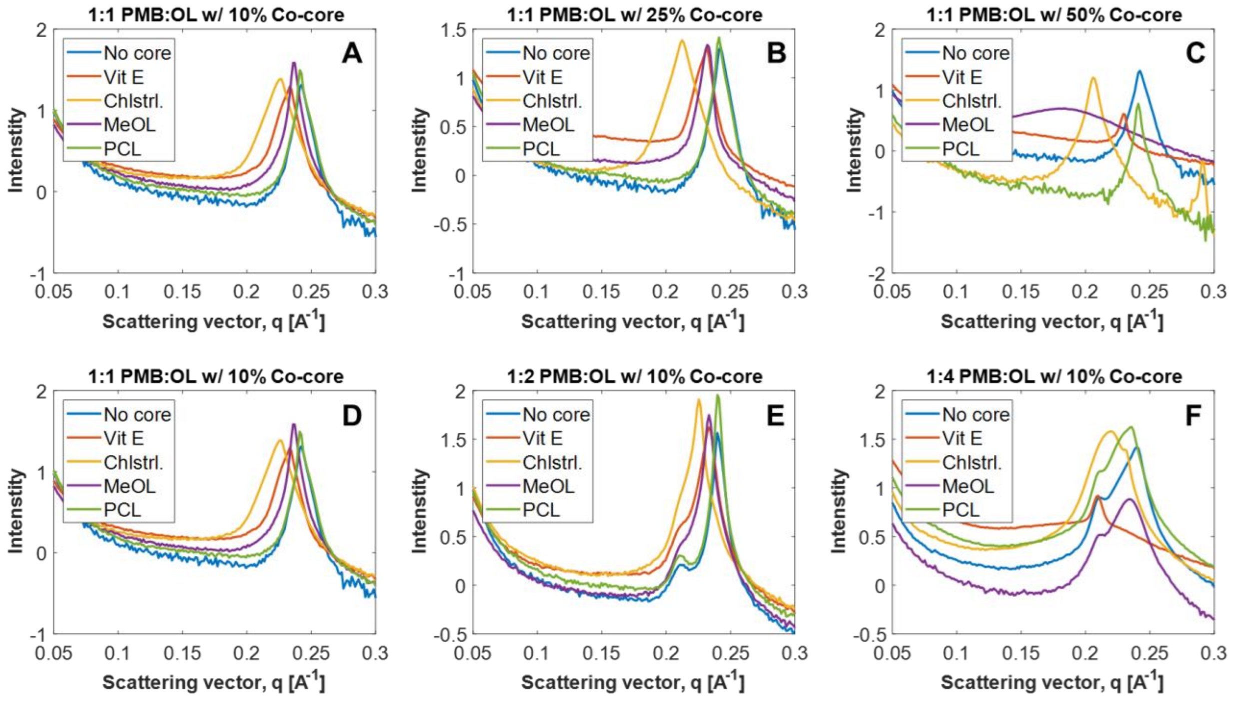

**Figure S.20:** PMB:OL LCNCs at a (A-B) Fixed 1:1 PMB:OL charge ratio investigation effects of increasing the co-core weight fraction and  (D) 1:1 vs (E) 1:2 vs (F) 1:4 charge ratio for a fixed 10% weight fraction co-core.
